# A Cortico-Striatal Circuit for Sound-Triggered Prediction of Reward Timing

**DOI:** 10.1101/2023.11.21.568134

**Authors:** Harini Suri, Karla Salgado-Puga, Yixuan Wang, Nayomie Allen, Kaitlynn Lane, Kyra Granroth, Alberto Olivei, Nathanial Nass, Gideon Rothschild

## Abstract

A crucial aspect of auditory perception is the ability to use sound cues to predict future events and to time actions accordingly. For example, distinct smartphone notification sounds reflect a call that needs to be answered within a few seconds, or a text that can be read later; the sound of an approaching vehicle signals when it is safe to cross the street. Other animals similarly use sounds to plan, time and execute behaviors such as hunting, evading predation and tending to offspring. However, the neural mechanisms that underlie sound-guided prediction of upcoming salient event timing are not well understood. To address this gap, we employed an appetitive sound-triggered reward time prediction behavior in head-fixed mice. We find that mice trained on this task reliably estimate the time from a sound cue to upcoming reward on the scale of a few seconds, as demonstrated by learning-dependent well-timed increases in reward-predictive licking. Moreover, mice showed a dramatic impairment in their ability to use sound to predict delayed reward when the auditory cortex was inactivated, demonstrating its causal involvement. To identify the neurophysiological signatures of auditory cortical reward-timing prediction, we recorded local field potentials during learning and performance of this behavior and found that the magnitude of auditory cortical responses to the sound prospectively encoded the duration of the anticipated sound-reward time interval. Next, we explored how and where these sound-triggered time interval prediction signals propagate from the auditory cortex to time and initiate consequent action. We targeted the monosynaptic projections from the auditory cortex to the posterior striatum and found that chemogenetic inactivation of these projections impairs animal’s ability to predict sound-triggered delayed reward. Simultaneous neural recordings in the auditory cortex and posterior striatum during task performance revealed coordination of neural activity across these regions during the sound cue predicting the time interval to reward. Collectively, our findings identify an auditory cortical-striatal circuit supporting sound-triggered timing-prediction behaviors.

## Introduction

In everyday life, sounds often predict forthcoming events, allowing for planning and execution of appropriate behavioral responses. Consider, for instance, the confidence with which we step out of an elevator’s open doors a few seconds after its chime, even without looking up from our phone. Or how the conclusion of a friend’s sentence determines the opportune moment for our response (Benichov et al., 2016; Levinson, 2016; Stivers et al., 2009). Similarly, we rely on distinct phone notification sounds to determine whether it is a call that we need to answer within a few seconds, or a text message that we can read a bit later. Likewise, animals rely on sound cues to gauge how swiftly they should vocalize in response to a conspecific call (Benichov & Vallentin, 2020), evade a predator (Z. Li et al., 2021), hone in on prey (Surlykke & Moss, 2000) or approach a needy offspring (Dunlap et al., 2020; Ehret, 2005). These and other examples in everyday life require humans and other animals to utilize sounds to predict and precisely time subsequent salient events and to initiate appropriate behavioral responses within the scale of seconds (Mazzucato, 2022).

The ability to use sounds to predict when future events will occur and consequently when to initiate appropriate action relies on a number of neural processing stages (Bueti, 2011; Buhusi & Meck, 2005; Wiener, Matell, et al., 2011). First, the sound must be detected, processed, and recognized. Second, the predicted amount of time from the sound to the future event is evaluated. And finally, action is initiated when the elapsed time matches the anticipated appropriate time to act. Neural signatures underlying the first stage of this process, namely sound processing and recognition, have been extensively identified in the auditory pathway, and in particular in the auditory cortex (For example, Suri & Rothschild, 2022; Bernal & Ardila, 2016; Geissler & Ehret, 2004; Jasmin et al., 2019; King et al., 2018; King & Schnupp, 2007; Read et al., 2002; Zatorre et al., 2002). Considerably less is known about the second stage, and specifically where and how the brain encodes the predicted amount of time from the sound to a future event. Traditional models of time perception have suggested the existence of a “centralized clock” in the brain (also referred to as the “Internal clock model”) (Hinton & Meck, 1997; Leow & Grahn, 2014; Treisman, 1963; Wearden, 2005). According to this model, the role of sensory regions is to detect the relevant sensory stimulus and communicate this information to higher-order centralized-clock brain regions, where a continuous representation of elapsed time is maintained (Buhusi & Meck, 2005; Hinton & Meck, 1997; Leow & Grahn, 2014; Treisman, 1963). This model proposes that the centralized clock is similarly able to estimate time based on cues from varying modalities, arriving via distinct pathways from the various sensory regions and hence, is “amodal” (Bueti, 2011; Wiener, Matell, et al., 2011). Different studies have implicated a number of brain regions as hosting such centralized clocks, including the nucleus accumbens (Kurti & Matell, 2011), caudoputamen (Matell et al., 2003), ventral tegmental area (Fiorillo et al., 2008), substantia nigra compacta (Meck, 2006), dorsal striatum (Jones & Jahanshahi, 2011; Meck, 2006; Wiener, Lohoff, et al., 2011) and the medial prefrontal cortex (Jones & Jahanshahi, 2011; Ning et al., 2022; Tunes et al., 2022).

However, the universality of this model has been challenged by studies showing that in addition to a centralized clock, there exist sensory-specific timing mechanisms, and that these are located within early sensory cortical regions (Bueti, 2011; Buhusi & Meck, 2005; Wiener et al., 2011). Early support for this suggestion came from studies demonstrating that the ability to estimate time from a sensory cue depends on the modality of that cue (For example - Hussain Shuler and Bear, 2006; Bueti et al., 2008). Furthermore, recent studies have identified signatures of time estimation within sensory cortices. For example, in vivo neural recordings in the primary visual cortex of rodents show various neural response forms which represent the time interval between the visual stimulus and the anticipated reward (Chubykin et al., 2013; Hussain Shuler & Bear, 2006; Namboodiri et al., 2015). A recent study further showed that this reward timing representation is modulated by an intracortical network of inhibitory interneurons in the visual cortex (Monk et al., 2020).

In the auditory pathway, a key candidate brain region for encoding sound-triggered timing is the auditory cortex (AC), due to its established role in behavior- and decision-making-dependent sound processing (Bathellier et al., 2012; Francis et al., 2018; Fritz et al., 2003, 2005; King & Schnupp, 2007; Lee & Rothschild, 2021; Nelken et al., 2014; Town et al., 2018; Vivaldo et al., 2023). Numerous studies have demonstrated retrospective coding of the degree to which a sound deviates from expectation in AC (Heilbron & Chait, 2018; Khouri & Nelken, 2015; Rubin et al., 2016; Taaseh et al., 2011; Ulanovsky et al., 2003, 2004; Yaron et al., 2012). However, much less is known about the existence of prospective coding of anticipated time from a sound to a subsequent event. In one study, auditory cortical responses to a tone series varied depending on whether a subsequent target sound was expected early (300-450ms) or late (1300-1500ms) within the series (Jaramillo & Zador, 2011). However, this study did not test whether AC encodes expectation of non-auditory cues or whether a continuous representation of predicted time is encoded. In a recent study, mice were trained on a self-paced action timing task, in which lever pressing caused reward delivery after 30 seconds. Optogenetic stimulation and inactivation of the secondary AC implicated its responses to the sound of lever press as being causally involved in timing reward-preparatory action (Cook et al., 2022). However, this study did not test whether and how AC is involved in encoding varying sound-reward time intervals, in particular on the scale of seconds. Thus, it remains unclear whether and how AC is involved in sound-triggered predictive timing of future salient events on the timescale of seconds.

The final step of sound-dependent prediction of imminent events requires initiation of appropriate action once the anticipated time from sound to action has elapsed. To carry this out, the neural information assimilated in the first two stages of this process needs to be sent to downstream brain regions to induce consequent action timing and initiation. Previous studies have implicated a number of brain regions that mediate sound-triggered action initiation, including the striatum (Guo et al., 2018; Matell et al., 2003; Tunes et al., 2022), the medial prefrontal cortex (Tunes et al., 2022; Yumoto et al., 2011) and the supplementary motor area (Mita et al., 2009; Wiener, Matell, et al., 2011). Anatomically, the most prominent candidate brain region to receive sound-triggered time keeping signals from AC and participate in converting this information to action, is the posterior tail of the dorsal striatum (hereafter referred to as the posterior striatum). The posterior striatum (pStr) receives monosynaptic projections from AC (Bertero et al., 2020; Huang et al., 2023; Znamenskiy & Zador, 2013), and these projections are known to be causally involved in auditory-guided decision making tasks (Znamenskiy & Zador, 2013) and auditory associative learning tasks (Huang et al., 2023). Moreover, pStr itself responds to auditory stimulation (Guo et al., 2018, 2019) and has been shown to play a key role in various stimulus-driven time keeping behaviors (Matell et al., 2003; Tunes et al., 2022). However, whether and how information about sound-triggered time interval estimation arriving from AC engages pStr remains unknown. To collectively address these gaps, we investigated the causal and functional role of the auditory cortical-striatal circuit in sound-triggered prediction of time to consequent reward.

## Results

### Mice use sound to predict reward time with 1-second temporal resolution

To investigate the role of auditory cortical-striatal neural mechanisms underlying sound-triggered reward timing prediction, we employed an appetitive sound-guided trace conditioning task in water-restricted mice. Following habituation to head fixation, 8 mice underwent training sessions of 150-200 trials each, in which each trial was initiated with a 1.5s long sound stimulus (composed of a sequence of three pure tones), followed by a fixed delay period and then delivery of water reward (Figure 1A). Trials were separated by inter-trial intervals which randomly varied between 2-6s in duration. All animals started behavioral training sessions with a fixed delay period of 1.5s between the sound cue termination and reward (“sound-reward time interval”). After 7-10 days of training, we introduced randomly interspersed 10-20% catch trials, in which the reward was withheld. Mice expressed learning of the sound-reward contingency by increasing their lick rate before the anticipated reward time (“Predictive licking”, Figure 1B, left) and by licking before and during the time of anticipated reward in the catch trials (Figure 1B, right).

**FIGURE 1:**
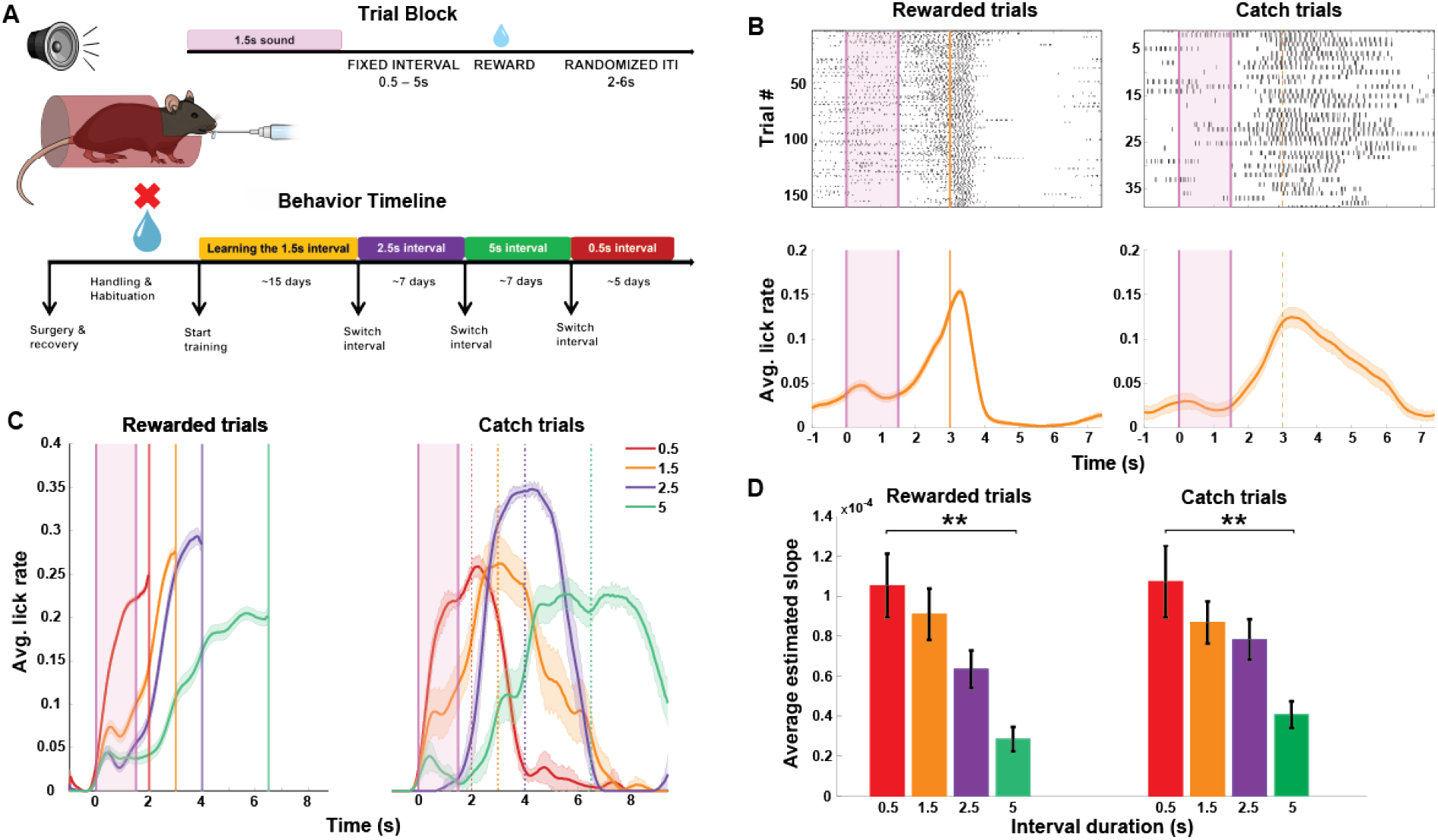
Mice predict reward timing using a sound cue. **A.** An illustration of the behavioral setup for sound-triggered reward time prediction task, components of a trial block and the experimental timeline for behavioral training. B. Top: Peri-sound lick raster of an example behavioral session from a trained animal performing on rewarded (left) and catch (right) trials within the session. Bottom: Average peri-sound lick rate response curve (solid line denotes mean, shaded area represents SEM across trials) for the example behavioral session above for rewarded (left) and catch (right) trials. Shaded pink region represents the 1.5s long sound period. Solid and dotted orange lines represent when reward was given in rewarded trials and expected in catch trials. Black ticks represent licks. C. Average peri-sound lick rate response curves (solid line denotes mean, shaded area represents SEM across trials) of an example animal trained to perform on the four different sound-reward intervals represented by different colors. Left: Rewarded trials; Right: Catch trials. Shaded pink region represents the 1.5s long sound period. Solid and dotted lines represent when reward was given in rewarded trials and expected in catch trials for each of the sound-reward interval. D. Average estimated slope of predictive licking response curves for each of the four sound-reward intervals across all animals (N = 8) for rewarded trials (left,**p=0.0064, Kruskal-Wallis test) and for catch trials (right, **p =0.0011, Kruskal-Wallis test). Error bars represent mean ± SEM across animals.

To determine whether predictive licking reflects a reliable estimation of anticipated reward time, the same mice then went on to train on the same paradigm but with different sound-reward time intervals. We posited that if mice can reliably estimate time on the scale of seconds and use these estimates to guide their behavior, the timing of predictive licking would vary with the duration of the sound-reward interval. To test this, once animals showed reliable performance on the task with the 1.5s sound-reward interval (as evidenced by reliable predictive licking) they relearned versions of this task with a 2.5s, 5s and 0.5s sound-reward time intervals, in this order (Figure 1A, Behavior Timeline). For each animal, we compared the slopes of their predictive licking curves on their “best” behavior days of each of these sound-reward time intervals (see Methods). We computed the slopes of the predictive licking curves for rewarded and catch trials separately (see Methods). We found that the predictive licking slopes varied with the duration of the sound-reward interval (Figure 1C). Across animals, the average slope of the predictive licking curves significantly varied as a function of the sound-reward interval duration, with the shortest interval duration (0.5s) inducing the steepest slope of predictive licking, followed by 1.5s, 2.5s and 5s (Figure 1D, Kruskal-Wallis test for multiple comparisons, p = 0.0011 for rewarded trials, p = 0.0064 for catch trials). We also quantified the precision of predictive licking for each of the sound-reward intervals by computing the full width at half maxima of the predictive licking curve for catch trials and found that mice are able to predict the shorter intervals with higher precision compared to the longer intervals (Supplementary Figure 1, Kruskal-Wallis test for multiple comparisons, p = 3.85×10^-6^). These results show that mice can estimate time intervals on the scale of 0.5-5s with at least 1-s temporal resolution and use these estimates to predict the time of expected reward following a sound.

### The auditory cortex is required for sound-triggered delayed reward prediction

The auditory cortex (AC) plays an important role in predictive and behavior-dependent sound processing (Bathellier et al., 2012; Francis et al., 2018; Heilbron & Chait, 2018; Kuchibhotla & Bathellier, 2018; Kuchibhotla et al., 2017; J. Li et al., 2017; Okada et al., 2018). Since our task requires mice to use the sound to time their behavior, we hypothesized that the AC is necessary for successful performance of this task. To test this hypothesis, we measured the influence of bilateral AC inactivation using the GABA-A receptor agonist muscimol on trained animals’ ability to predict reward at a 1.5s sound-reward time interval. Muscimol expression was histologically validated at the end of each experiment and only animals with selective targeting in AC were included (Figure 2A). AC inactivation resulted in an overall reduction in the ability to predictively lick for reward as compared to infusion of inert PBS as a control (Figure 2B). An impaired ability to predictively lick for reward following muscimol infusion was consistently observed in all our animals (N=8, p<0.05 for each mouse, Wilcoxon rank-sum test). To rule out the possibility that reduced predictive licking reflects a simple impairment in sound processing, we trained another cohort of mice on a variation of the task, in which reward was delivered immediately following sound onset (No-Delay Task). Mice trained on the No-Delay task showed evidence for sound-reward association by consistently licking in response to the sound during catch trials, when no reward was delivered (Figure 2C, right). Interestingly, this form of predictive licking following the No-Delay sound-reward association was unaffected by AC inactivation (N=8, p>0.05 for each mouse, Wilcoxon rank-sum test; Figure 2C). To directly compare the influence of AC inactivation on prediction of delayed and immediate reward, we calculated the log of the ratio of predictive licking in PBS and muscimol (“log PLR”) for each of the task versions. Thus, larger log PLR values indicate a greater influence of AC inactivation on predictive licking. Using this metric, we found that AC inactivation had a significantly more detrimental effect on sound-guided prediction of 1.5 s-delayed reward than on immediate reward (p=0.000156, Wilcoxon rank-sum test, Figure 2D). These findings demonstrate that the AC is causally involved in sound-triggered time-delayed reward prediction.

**FIGURE 2:**
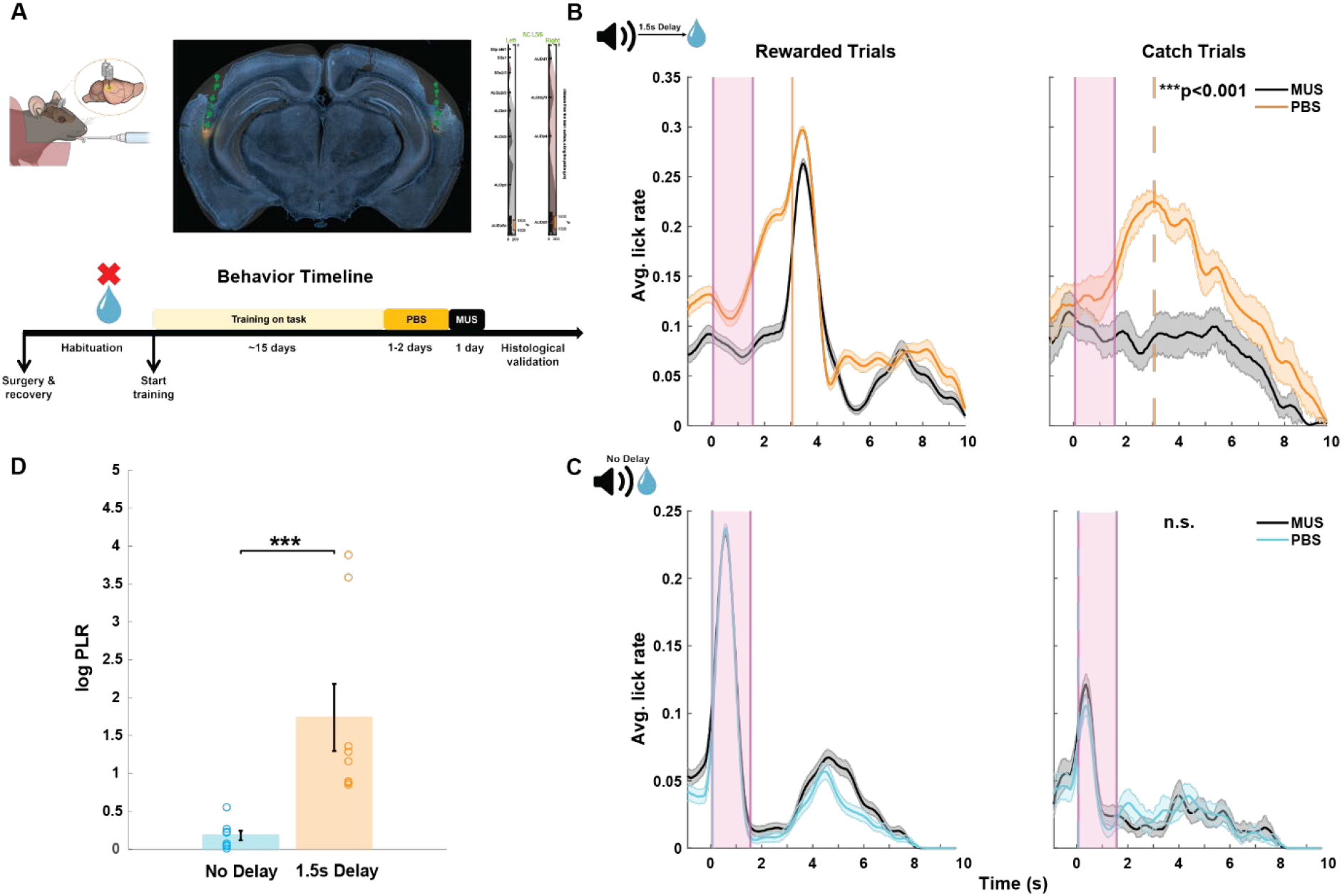
AC is required for sound-triggered delayed reward prediction. A. Top: An illustration of the cannula implanted in AC for muscimol infusion. Right: Histological verification of bilateral muscimol infusion into AC. The brain slice acquired from an example animal is overlaid with the corresponding coronal section from the Allen Mouse Brain Atlas (see Methods). The cannula tracks are indicated by the green dotted line on the brain slice and the site of muscimol infusion is seen in orange. Markers on the right indicate the depth at which muscimol was infused in the left and right hemispheres. Scale bar: 1000µm. Bottom: Behavioral timeline for mice that underwent training on the 1.5s Delay task. B. Average peri-sound lick rate response curves (solid line denotes mean, shaded area represents SEM across trials) of an example animal trained to predict reward at 1.5s sound-reward interval when infused with PBS (orange) and muscimol (MUS, black) in AC for rewarded trials (left) and catch trials (right). Shaded pink region represents the sound period. Solid and dotted orange lines represent when reward was given in rewarded trials and expected in catch trials. ***p=0.0007 (Wilcoxon rank-sum test) denotes the significant difference in predictive licking compared to baseline for catch trials on PBS and MUS conditions. C. Average peri-sound lick rate response curves (solid line denotes mean, shaded area represents SEM across trials) of an example animal trained on the No-Delay task when infused with PBS (light blue) and muscimol (MUS, black) in AC for rewarded trials (left) and catch trials (right). Shaded pink region represents the 1.5s long sound period. Solid and dotted blue lines represent when reward was given in rewarded trials and expected in catch trials. n.s. (p=0.939, Wilcoxon rank-sum test) denotes the not significant difference in predictive licking between catch trials on PBS and MUS in the No-Delay task. D. Significant difference in average log predictive licking ratio (log PLR) between No-Delay (N = 8) and 1.5s Delay (N = 8) cohorts (***p =0.000156, Wilcoxon rank-sum test). Error bars represent mean ± SEM across animals. Open circles indicate the PLR for each animal in the No-Delay (blue) and 1.5s Delay cohorts.

### Mice predict reward timing from sound onset

Having established that mice can use the sound cue to predict the timing of reward across varying sound-reward time intervals and that this behavior causally involves the auditory cortex, we next asked whether this behavior reflects time estimation from sound onset or from sound termination (Figure 3A). To this end, we compared the predictive lick pattern under the standard paradigm of a 1.5s-long sound and 1.5s-long sound-reward interval, to a similar paradigm in which the sound duration was cropped by 0.5s, to a duration of 1s. We argued that if mice use sound onset to predict reward timing, their predictive lick pattern would be unaffected by a shorter sound duration, whereas if mice use sound termination to predict reward timing, predictive licking will start earlier in the trials with the shorter sound cue. To test this, we trained a cohort of 8 mice to predict reward at a 1.5s sound-reward interval and then introduced them to a training session with 35% short sound trials (Figure 3A). All our animals showed similar behavioral responses on short sound and standard sound trials, with almost identical predictive licking curves (Figure 3B). Across animals, the slopes for the predictive licking curves were not significantly different between the two types of trials (Figure 3C, Wilcoxon rank-sum test, p = 0.96), and the average log PLR for standard sound to short sound trials was not significantly different from 0 (See Methods, Figure 3D, Wilcoxon signed rank test compared against 0, p = 0.12). These results suggest that mice primarily use the sound onset to estimate the amount of time from sound to reward.

**FIGURE 3:**
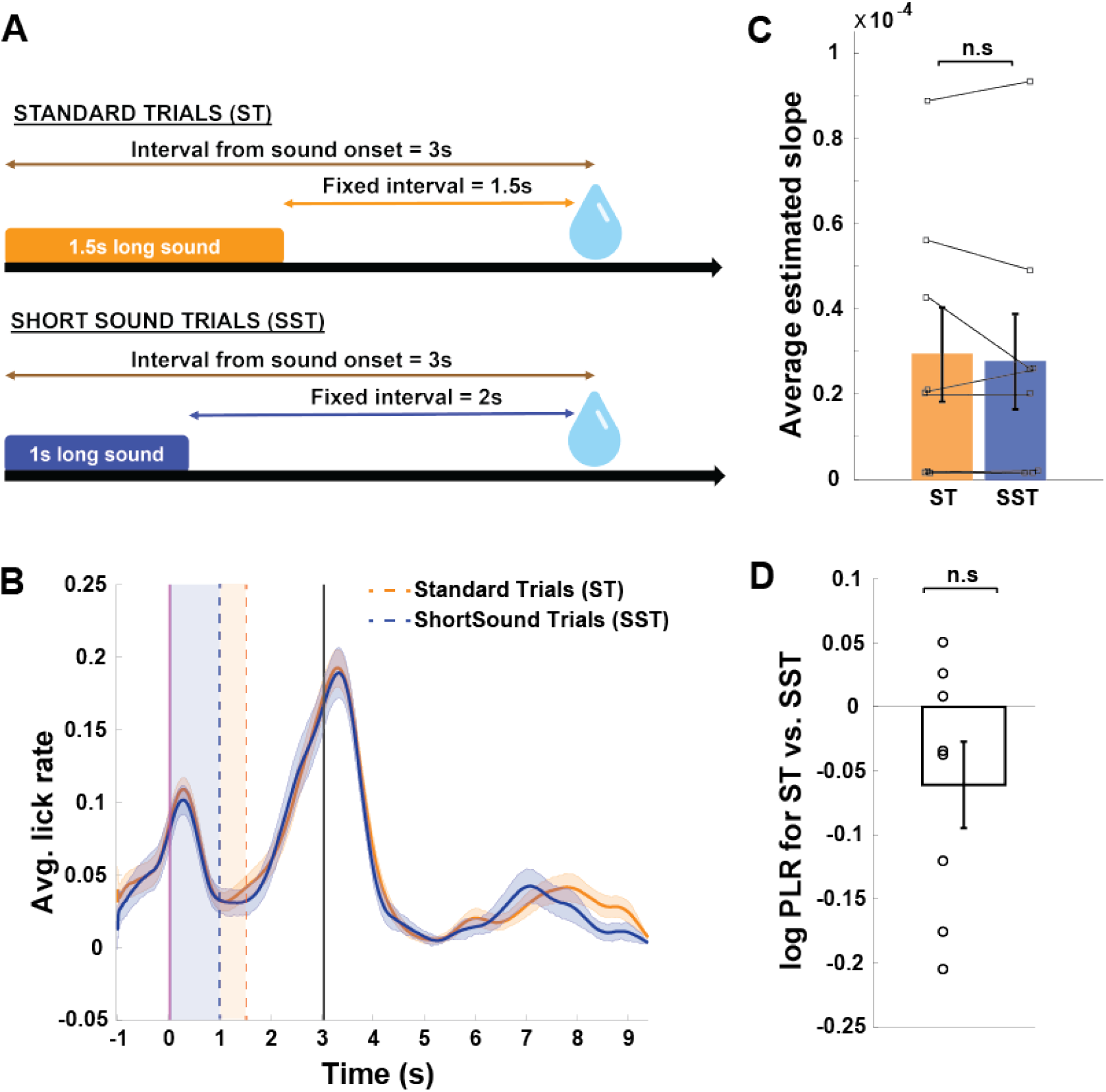
Mice predict reward timing from sound onset. A. An illustration of the trial designs. B. Average peri-sound lick rate response curves (solid line denotes mean, shaded area represents SEM across trials) of an example animal trained on the Standard Trials (ST, orange) and Short Sound Trials (SST, navy blue). Dashed lines indicate the sound termination times for ST (orange) and SST (blue) trials. Solid black line represents when reward was given. Comparison of the predictive licking between ST and SST trials compared to baseline yields p = 0.9 (Wilcoxon rank sum test). C. Average estimated slope of the predictive licking curves for ST (orange) and SST (navy blue) across animals (N = 8, p = 96, Wilcoxon rank-sum test). Error bars represent mean ± SEM across animals. Lines connecting filled squares represent the estimate slope values for each animal in ST and SST. D. Average log predictive licking ratio (log PLR for SST) across animals is not significantly different from 0 (N = 8, p = 0.12, Wilcoxon signed rank test compared against 0). Error bars represent mean ± SEM across animals. Open circles represent the PLR for each animal.

### AC sound responses encode predicted time-to-reward

A prominent model for the neural processing underlying sound-triggered time predictions suggests that following sound coding in the auditory pathway, downstream brain regions encode the anticipated time interval from the sound to subsequent events (Buhusi & Meck, 2005; Hinton & Meck, 1997; Leow & Grahn, 2014; Treisman, 1963). Alternatively, given the established role of the AC in predictive coding, the predicted time from sound to reward may already be reflected in the AC response to the sound itself. To test this possibility, we recorded local field potential (LFP) activity in the AC of the right hemisphere in 8 mice, as they were trained to predict timed reward using sound at the four different sound-reward intervals in the order described previously. We histologically verified the position of these electrodes at the end of the experiments (Figure 4A). Similar to our findings in the first cohort (Figure 1), animals in this cohort also changed the rate of their predictive licking curves according to the duration of the sound-reward intervals, with the slope increasing with increased interval durations for both rewarded and catch trials (Figure 4B and C, p = 0.0000544 for rewarded trials and p = 0.00012 for catch trials; Kruskal-Wallis test for group differences). We similarly identified the best behavior performance day of each sound-reward interval for analysis and excluded trials in which the animal moved or in which the animal licked in the 200 ms period from sound onset to avoid a contribution of motor activity to the sound response magnitude (Schneider et al., 2014; Vivaldo et al., 2023; Whitton et al., 2014). We then quantified the magnitude of the AC responses to the sound for each of the interval durations per animal. As our behavioral results showed that mice use the sound onset to estimate the time to reward, we focused on the responses to sound onset. Interestingly, we found that AC responses (to the same sound) increased in magnitude with increasing sound-reward intervals (Figure 4D). Across animals, the average normalized sound response magnitude was significantly different across interval durations (p=1.9×10^-35^, Kruskal-Wallis test), with the response magnitude increasing as a function of the time interval from sound to reward (Figure 4E). We also quantified the magnitude of responses to the sound offset for each of the interval durations per animal and found that these offset responses did not significantly change in magnitude across sound-reward intervals (p = 0.263, Kruskal-Wallis test, Figure 4F and G). Together, these results indicate that AC neural responses to the sound onset encodes, beyond the sound itself, the predicted time from sound to reward.

**FIGURE 4:**
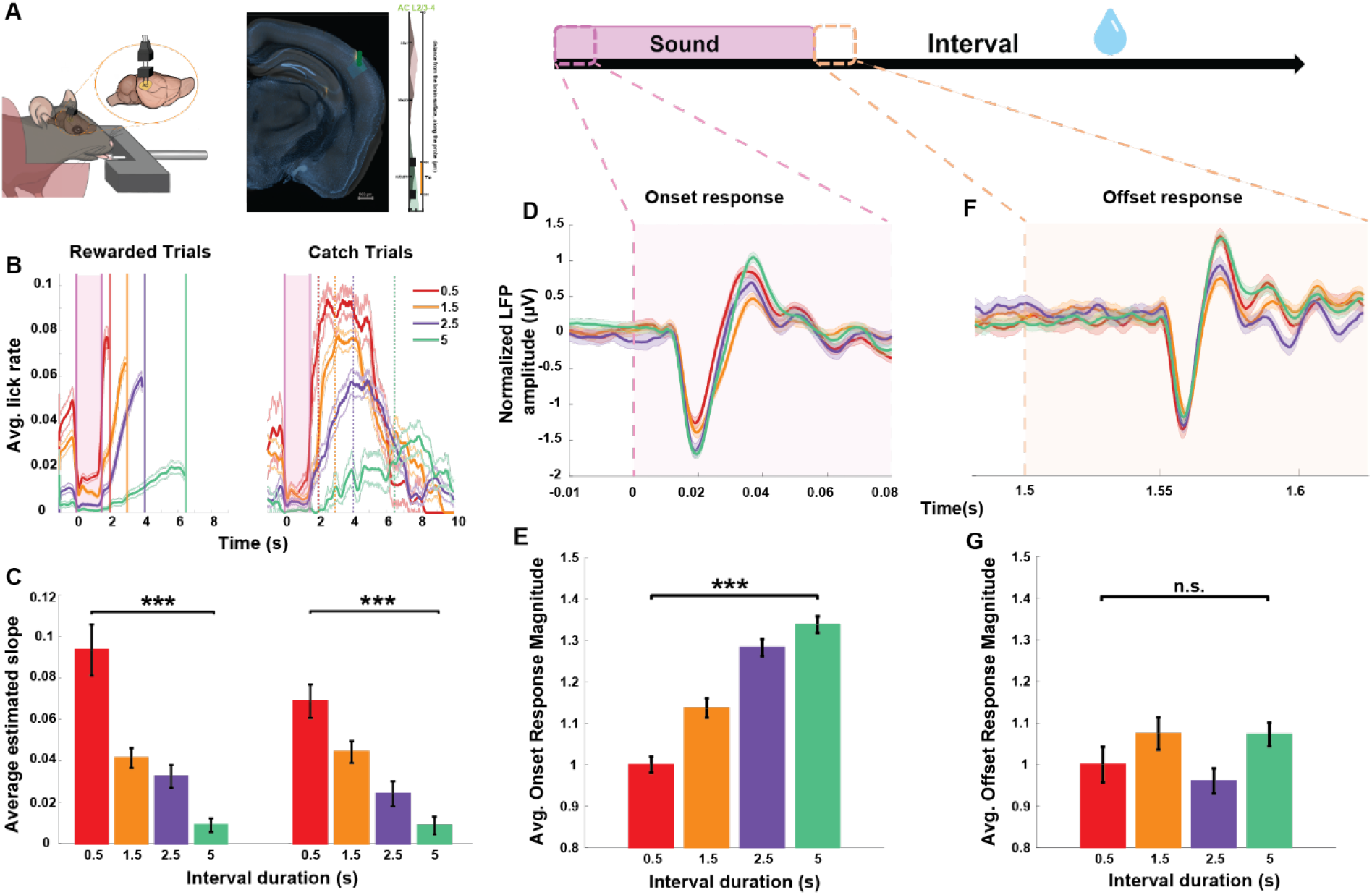
Auditory cortical sound responses encode predicted time to reward. A. Left: An illustration of the electrodes implanted in right AC. Right: Histological verification of electrode position in AC of the right hemisphere. The brain slice acquired from an example animal is overlaid with the corresponding coronal section from the Allen Mouse Brain Atlas (see Methods). The electrode track is indicated by the green dotted line on the brain slice and the markers on the right indicate the depth at which the electrode was implanted in the right hemisphere. Scale bar: 500µm B. Average peri-sound lick rate response curves (solid line denotes mean, shaded area represents SEM across trials) of an example animal trained to perform on the four different sound-reward intervals represented by different colors. Left: Rewarded trials; Right: Catch trials. Shaded pink region represents the 1.5s long sound period. Solid and dotted lines represent when reward was given in rewarded trials and expected in catch trials for each of the sound-reward interval. C. Average estimated slope of predictive licking response curves for each of the four sound-reward intervals across all animals (N = 8) for rewarded trials (left,***p=0.000054, Kruskal-Wallis test) and for catch trials (right, ***p =0.00012, Kruskal-Wallis test). Error bars represent mean ± SEM across animals. D. Normalized average AC LFP responses to the sound onset from an example animal trained on the four different sound-reward intervals represented by the different colors (solid line denotes mean, shaded area represents SEM across no-lick trials). The shaded pink region represents the period from sound onset as shown in the illustration above. E. Average onset response magnitude computed across animals (N = 8) for each of the sound-reward intervals. Error bars represent mean ± SEM across animals. Comparison across sound-reward intervals yields ***p = 1.9×10^-35^ (Kruskal-Wallis test). F. Normalized average AC LFP responses to the sound offset from an example animal trained on the four different sound-reward intervals represented by the different colors (solid line denotes mean, shaded area represents SEM across no-lick trials). The shaded yellow region represents the period from sound onset as shown in the illustration above. G. Average offset response magnitude computed across animals (N = 8) for each of the sound-reward intervals. Error bars represent mean ± SEM across animals. Comparison across sound-reward intervals yields p = 0.263 (Kruskal-Wallis test).

### Auditory cortical projections to posterior striatum are causally involved in sound-guided prediction of delayed reward

For successful sound-triggered reward timing prediction, animals need to translate their prediction of reward time into motor action, which in the current task is licking. While our data suggests that the AC is involved in sound-triggered reward timing prediction, we hypothesized that this behavior and its translation into motor action would further depend on the posterior striatum (pStr). The pStr is a key candidate brain region to support this function, as it receives strong monosynaptic projections from the AC (Bertero et al., 2020; Huang et al., 2023; Znamenskiy & Zador, 2013) as well as from the thalamic medial geniculate body (Huerta-Ocampo et al., 2014; LeDoux et al., 1991; Smeal et al., 2008) and is involved in sound processing and sound-guided behaviors (Huang et al., 2023; Znamenskiy & Zador, 2013). Moreover, pStr itself is known to be involved in appetitive auditory frequency discrimination tasks (Guo et al., 2018, 2019). Hence, we investigated the role of auditory cortical projections to the pStr in sound-triggered reward time prediction.

To address this, we chemogenetically inactivated the projections from AC to pStr in animals trained to predict timed reward using a sound cue. Using a dual virus approach, we bilaterally injected and expressed a Cre-dependent DREADD virus, AAV-hSyn-DIO-hM4D(Gi)-mCherry, in the AC of mice in the experimental group, (or AAV-hSyn-DIO-mCherry in the control group), and a retrograde-CRE virus (pENN/AAVrg-hSyn-Cre-WPRE-hGH) in the pStr. This approach allowed us to specifically target the projections from the AC to pStr for chemogenetic inactivation. The expression of these viruses was histologically validated at the end of the experiments and only those with targeted virus expression in AC cell bodies and pStr axonal projections were included (Figure 5B).

**FIGURE 5:**
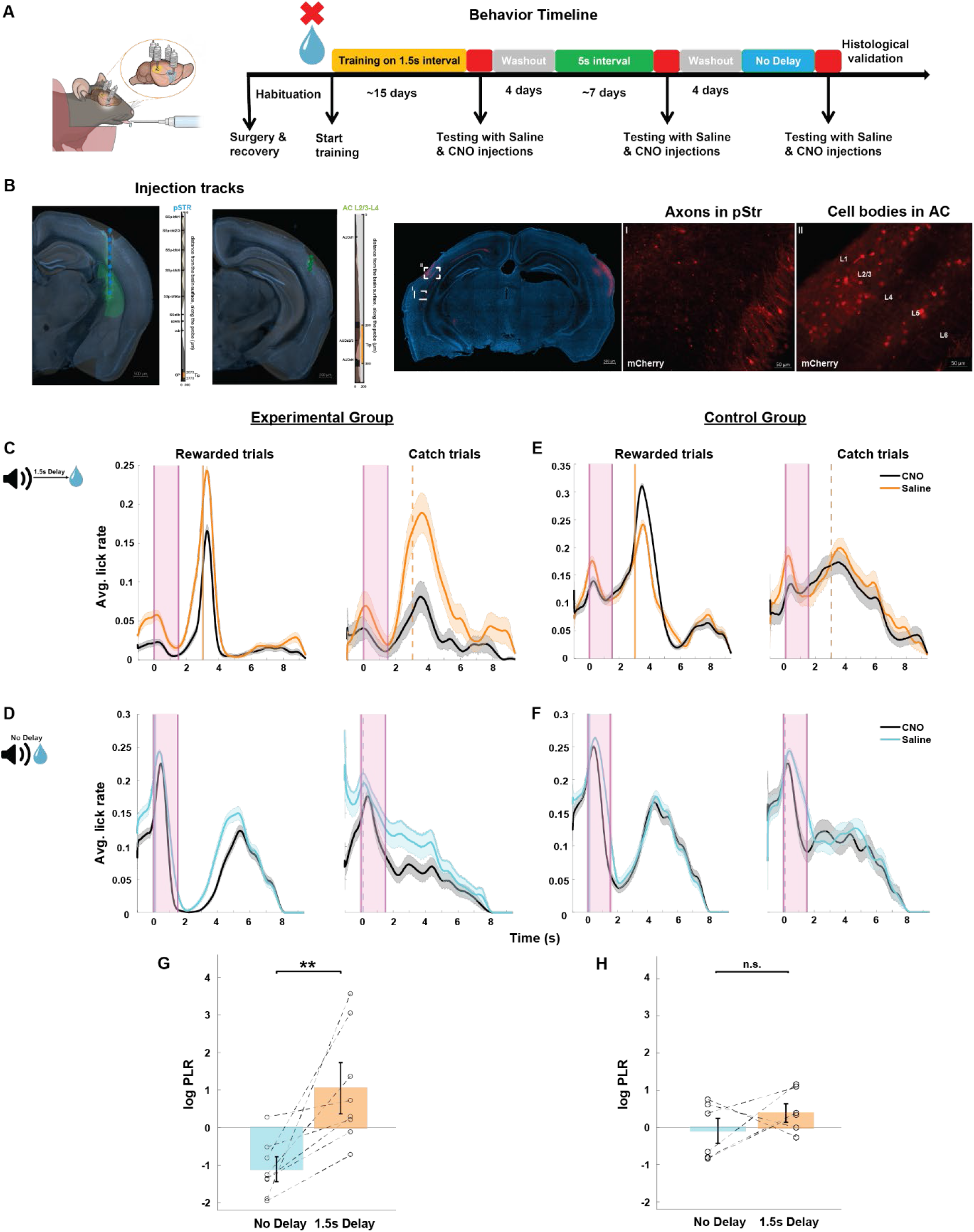
AC to pStr projections are causally involved in sound-triggered delayed reward prediction. A. Left: An illustration of cannulas implanted in bilateral AC and pStr for virus injections. Right: Behavioral timeline for chemogenetic inactivation experiments. B. Histological verification of selective virus expression in the projections from AC to pStr. A brain slice acquired from an example animal is overlaid with the corresponding coronal section from the Allen Mouse Brain Atlas for both and AC and pStr (see Methods). Left: Virus injection tracks in pStr and AC denoted by green dotted line on the brain slice and the markers on the right indicate the depth at which the electrode was implanted in the right hemisphere. Scale bar: 500µm Right: Magnified images of DREADD virus expression in pStr axons (I) and AC cell bodies (II). Scale bar: 50µm C. to F. Average peri-sound lick rate response curves (solid line denotes mean, shaded area represents SEM across trials) of an example animal trained to predict reward at either 1.5s sound-reward interval (C and E) or at no delay (D and F) when injected with saline (orange or light blue) and CNO (black) for rewarded trials (left) and catch trials (right). Shaded pink region represents the 1.5s long sound period. Solid and dotted orange or light blue lines represent when reward was given in rewarded trials and expected in catch trials. Left column represents animals from the experimental group and right column represents animals from the control group. G. and H. Average log Predictive Licking Ratio (log PLR) across animals trained on the 1.5s Delay and No-Delay tasks in the experimental group (left, N = 8) and in the control group (right, N=6). Lines connecting the circles represent the log PLR for each animal when trained on the 1.5s Delay and No-Delay tasks. The average log PLR for experimental group animals was significantly higher for 1.5s Delay task than No-Delay task (**p = 0.0019, Wilcoxon rank-sum test) and was not significantly different for the control group animals (p = 0.366, Wilcoxon rank-sum test).

Mice were trained to reliably predict sound-guided reward at a 1.5s sound-reward interval. They were then injected with saline (i.p.) as a control and their behavioral performance was recorded 30 minutes after the injection. 24 hrs later, we injected them with CNO (5mg/kg, i.p.) and after 30 min again recorded their behavioral performance (Figure 5A). Chemogenetic silencing of the AC-pStr projections in the mice of the experimental group significantly reduced their ability to predict reward at 1.5s (Figure 5C). We observed this significant effect in all mice in the experimental group (p<0.05 for each mouse, Wilcoxon rank-sum test). In contrast, none of the mice in the control group showed a significant change in predictive licking following CNO injection compared to their performance with saline injection (Figure 5F, p>0.05 for each mouse, Wilcoxon rank-sum test). These findings demonstrate that the AC-pStr projection is necessary for sound-triggered prediction of delayed reward.

Following a 4-day CNO-washout period, we trained a subset of these mice in the experimental and control groups to use the same sound cue to predict reward at a 5s interval from sound (Figure 5A). We chemogenetically inactivated the AC-pStr projections again and found that all mice in the experimental group showed poor reward prediction at 5s interval following CNO injection compared to their performance with saline injection on the previous day (Supplementary Figure 2A, p<0.05 for each mouse, Wilcoxon rank-sum test). In contrast, mice in the control group did not show an impairment in their ability to predict timed reward at 5s from saline to CNO behavioral sessions (Supplementary Figure 2B, p>0.05 for each mouse, Wilcoxon rank-sum test).

To verify that chemogenetic inactivation of AC-pStr projections does not impair animals’ ability to process sounds and form sound-reward associations, we trained all mice in the experimental and control groups on the No-Delay task following another 4-day CNO washout period and followed the same manipulation protocol above (Figure 5A). Mice in both these groups did not show an impairment in their ability to lick for reward immediately following sound in the catch trials (Figure 5D and G).

At a population-level, we quantified the effect of the AC-pStr projection inactivation on behavior using the log PLR for Saline to CNO days (see Methods), such that higher values indicate a reduction in animals’ ability to reliably predict the time-to-reward from sound. We found that the average log PLR was significantly higher for the 1.5 s sound-reward delay compared to the No-Delay (Figure 5E, p=0.0019, Wilcoxon rank-sum test). These ratios did not significantly differ between the 1.5s and 5s delays (Supplementary Figure 2C, p =0.808, Wilcoxon rank-sum test). In contrast, mice in the control group showed no significant difference in the average log PLR across No-Delay, 1.5s and 5s delays (Figure 5H, Supplementary Figure 2D, p=0.1774, Kruskal-Wallis test). These findings further establish the causal involvement of the AC-pStr projections in animals’ ability to predict delayed reward using a sound cue.

To investigate the functional consequence of inactivation of the AC-pStr pathway on pStr activity, in a subset of 3 animals from the experimental group, we acquired LFP responses in pStr as these animals trained on the 1.5s-Delay and No-Delay tasks. We filtered for no-lick trials and computed the average response magnitude of pStr responses at sound onset. pStr sound onset response magnitude significantly reduced in all the 3 animals from saline to CNO days for the 1.5s-Delay task (Supplementary Figure 3B and C, p<0.05 for each mouse, Wilcoxon rank-sum test). We did not observe a significant change in pStr onset response magnitude from saline to CNO days for the No-Delay task (Supplementary Figure 3D and E, p>0.05 for each mouse, Wilcoxon rank-sum test). This selective reduction in pStr LFP response magnitude following the chemogenetic inactivation of the AC-pStr projections in the 1.5s-Delay task provides neurophysiological corroboration of the causal involvement of these AC-pStr projections in prediction of reward timing using a sound cue. Together, these findings show that AC-pStr projections are a necessary neural pathway for sound-guided delayed reward prediction behavior.

### The posterior striatum is involved in sound-guided reward time prediction

We next asked whether the target region of these AC-pStr projections, pStr itself, is a necessary component of the neural circuitry supporting sound-guided reward prediction behavior. Mice were bilaterally implanted with cannulae into their pStr (Figure 6A). We compared their ability to predict reward at 1.5s with muscimol infusion to that with PBS infusion on a day prior and found that inactivation of pStr reduced animals’ ability to predict the time-delayed reward (Figure 6B, p<0.05, Wilcoxon rank-sum test). Similar to the previous inactivation experiments, we trained another cohort of mice on the No-Delay task and compared their behavioral performances following PBS and muscimol infusion into pStr. Inactivation of pStr using muscimol did not change animals’ behavior on the No-Delay task (Figure 6C, p>0.05, Wilcoxon rank-sum test), confirming that there was no impairment in animals’ ability to process sounds or lick for reward. Across animals, the average log PLR for PBS to muscimol days for the 1.5s Delay task was significantly higher than that of the No-Delay task (Figure 6D, p = 0.0079, Wilcoxon rank-sum test). Overall, these findings show that pStr is also required for successful sound-guided reward prediction behavior.

**FIGURE 6:**
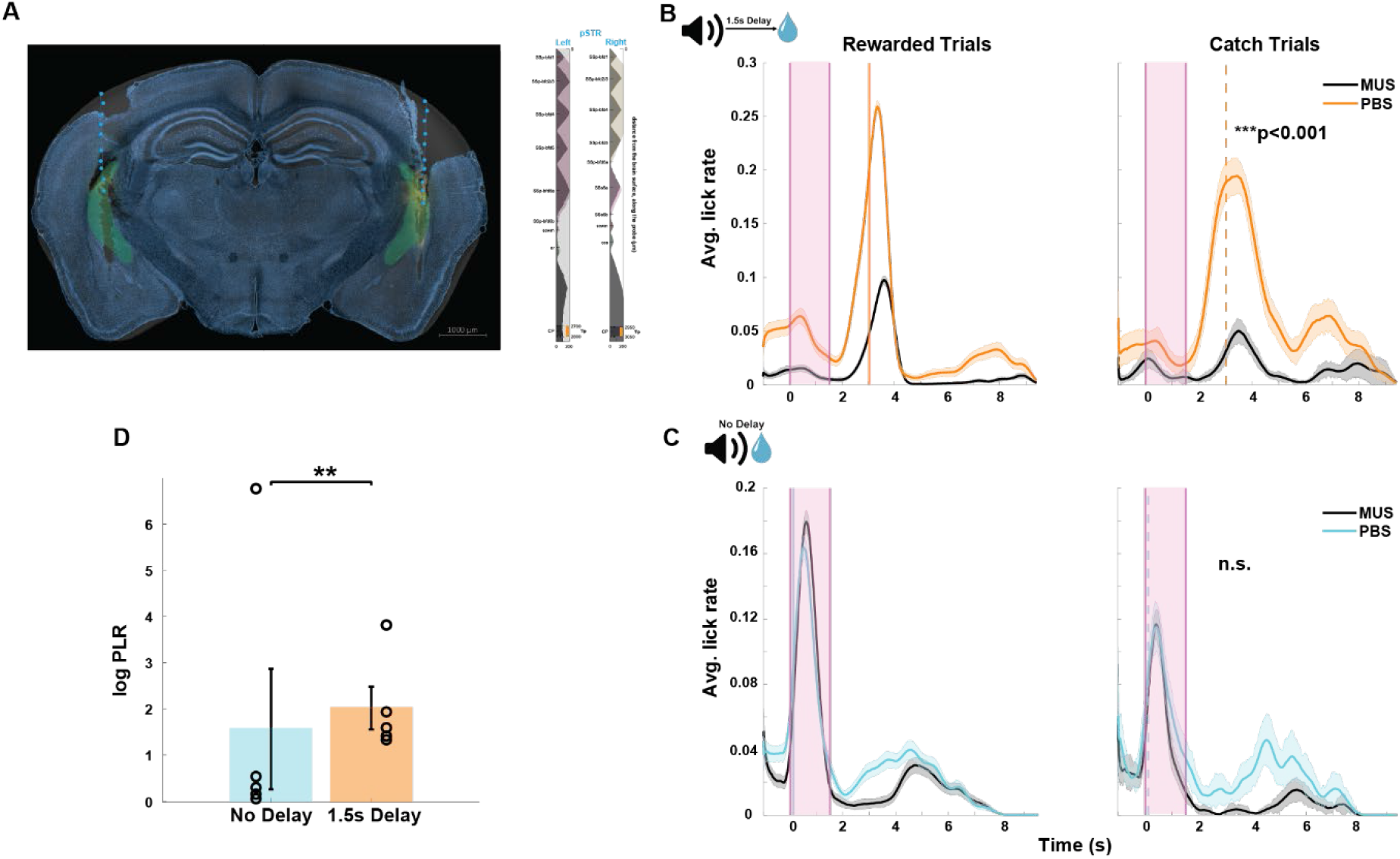
pStr is causally involved in sound-triggered delayed reward prediction. A. Histological verification of muscimol expression in bilateral pStr in a representative animal’s brain slice. The brain slice acquired from an example animal is overlaid with the corresponding coronal section from the Allen Mouse Brain Atlas (see Methods). The cannula tracks are indicated by the green dotted line on the brain slice and the site of muscimol infusion is seen in orange. Markers on the right indicate the depth at which muscimol was infused in the left and right hemispheres. Scale bar: 1000µm B. Average peri-sound lick rate response curves (solid line denotes mean, shaded area represents SEM across trials) of an example animal trained to predict reward at 1.5s sound-reward interval when infused with PBS (orange) and muscimol (MUS, black) in pStr for rewarded trials (left) and catch trials (right). Shaded pink region represents the 1.5s long sound period. Solid and dotted orange lines represent when reward was given in rewarded trials and expected in catch trials. ***p=0.000007(Wilcoxon rank-sum test) denotes the significant difference in predictive licking between catch trials on PBS and MUS. C. Average peri-sound lick rate response curves (solid line denotes mean, shaded area represents SEM across trials) of an example animal trained on the No-Delay task when infused with PBS (light blue) and muscimol (MUS, black) in pStr for rewarded trials (left) and catch trials (right). Shaded pink region represents the 1.5s long sound period. Solid and dotted blue lines represent when reward was given in rewarded trials and expected in catch trials. n.s. (p=0.3, Wilcoxon rank-sum test) denotes the not significant difference in predictive licking between catch trials on PBS and MUS in the No-Delay task. D. Significant difference in average log predictive licking ratio (log PLR) between No-Delay (N = 5) and 1.5s Delay (N = 5) cohorts (**p =0.0079, Wilcoxon rank-sum test). Error bars represent mean ± SEM across animals. Open circles indicate the PLR for each animal in the No-Delay (blue) and 1.5s Delay cohorts.

### Coordination of sound-evoked responses in AC and pStr during sound-triggered reward time prediction

To determine the activity patterns in pStr and to test whether the AC and pStr activity is coordinated during sound-triggered reward time prediction behavior, we simultaneously recorded LFP activity in AC and pStr as animals learnt to predict sound-triggered reward timing at the four sound-reward time intervals (Figure 7A). Like we did for analyzing AC responses, we eliminated trials with movement and included only trials with no licks in the first 200ms from sound onset for the best behavior days on each sound-reward interval. As expected from previous studies (Guo et al., 2018, 2019), we found robust sound responses in pStr, albeit of lower magnitude than in the AC. When we compared the average pStr sound responses across all four sound-reward intervals, we found that pStr responses also tended to increase with sound-reward interval duration (Figure 7B). However, at the population level, we noticed that while the pStr response magnitude was significantly different across sound-reward intervals (Figure 7C, p = 8.96×10^-6^, Kruskal-Wallis test), the pair-wise comparison of the response magnitude across all pairs of sound-reward intervals did not yield significant differences, unlike responses in the AC.

**FIGURE 7:**
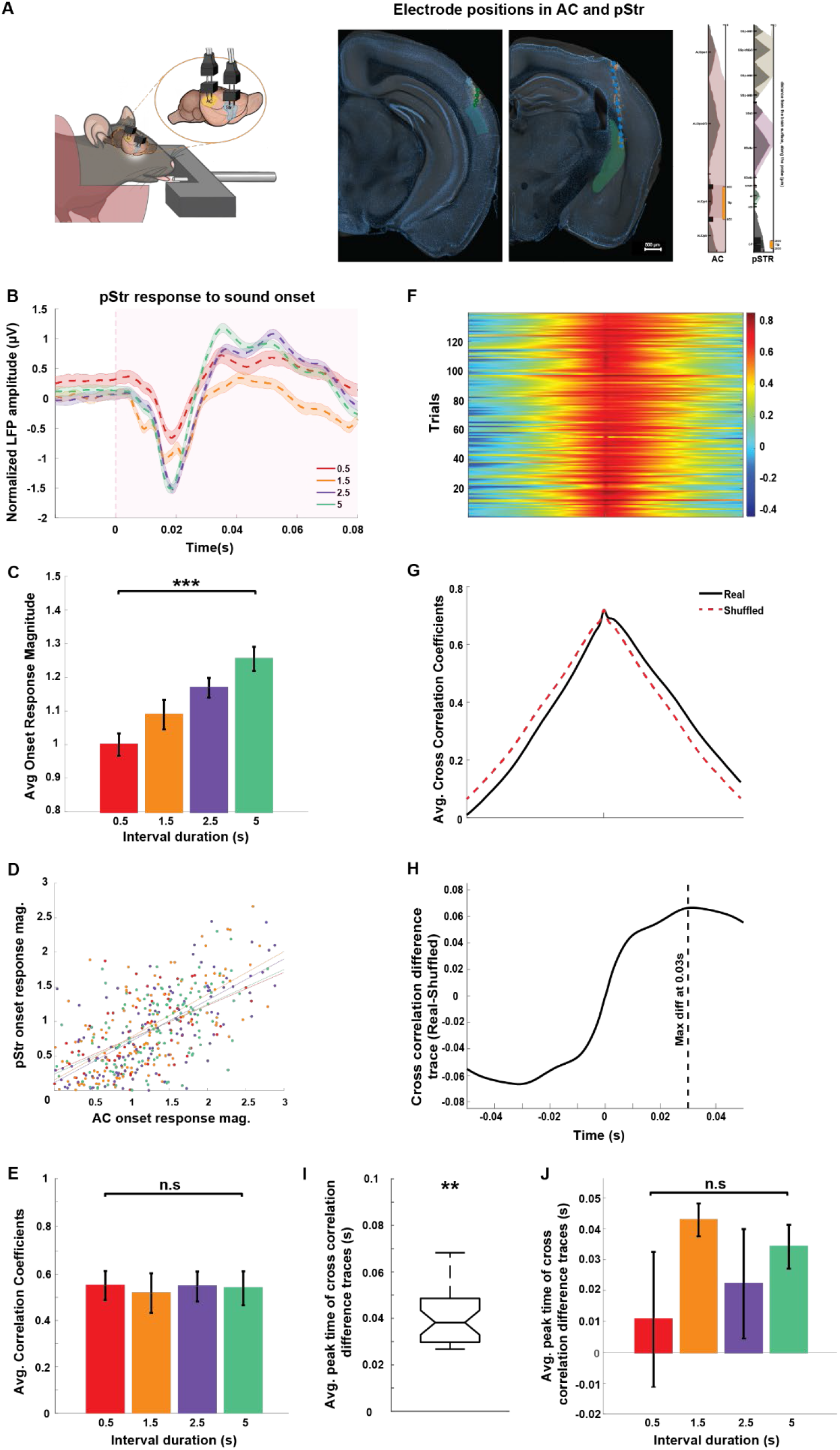
Coordination of LFP activity across AC and pStr during sound-triggered reward time prediction. A. Left: Illustration of electrode implanted in right AC and pStr. Right: Histological verification of electrode positions in AC (left) and pStr (right). The brain slice acquired from an example animal is overlaid with the corresponding coronal section from the Allen Mouse Brain Atlas (see Methods). The electrode tracks are indicated by the green dotted line on the brain slices and the markers in the middle indicate the depth at which the electrode was implanted in the right hemisphere. Scale bar: 500µm. B. Normalized average pStr LFP (solid line denotes mean, shaded area represents SEM across no-lick trials) recorded in response to the sound onset from an example animal trained on the four different sound-reward intervals (represented by the different colors). The shaded pink region represents the period from sound onset. C. Average pStr onset response magnitude computed across animals (N = 8) for each of the sound-reward intervals. Error bars represent mean ± SEM across animals. Comparison across sound-reward intervals yields ***p = 8.96×10^-6^ (Kruskal-Wallis test). D. Scatter plot of the trial-wise correlation between AC and pStr onset response magnitudes for each of the sound-reward intervals for an example animal. Filled circles of different colors represent the trial-wise onset response magnitudes and the solid lines represent the linear fits for each of the sound-reward intervals. Pearson correlation coefficients for each of the sound-reward intervals was positive and significant: 0.5s interval (red): r = 0.582, p<0.001; 1.5s interval (orange): r = 0.673, p<0.001; 2.5s interval (purple): r = 0.673, p<0.001; 5s interval (green): r = 0.561, p<0.001. E. The average correlation coefficients across animals across sound-reward intervals are not significantly different (p = 0.99, Kruskal-Wallis test) F. Heat map representing the trial-wise cross-correlation of AC and pStr sound-evoked LFP responses for an example animal trained on an 0.5s sound-reward interval behavioral session. Time axis is set such that it denotes AC leading on the positive side and pStr leading on the negative side. Color bar indicates the cross-correlation coefficients. G. Average cross correlation of AC and pStr sound evoked LFP responses across trials for the behavioral session shown above is indicated in black. The dotted red line shows the average of the shuffled cross correlation computed from randomized AC and pStr sound evoked LFP responses from the session. H. The solid black line shows the difference trace between the real and shuffled average cross correlation in panel G. This cross-correlation difference trace has a peak at 0.03s as indicated by the dotted black line. I. The average of the peak times of the cross-correlation difference traces computed across all animals and across all sound-reward intervals is significantly greater than 0s (**p= 0.0084, Wilcoxon signed rank test). J. Average peak times of the cross-correlation difference traces across all animals was not significantly different across sound-reward intervals (p = 0.94, Kruskal-Wallis test).

Next, we asked whether neural activity across the two brain regions was coordinated during sound-guided reward time prediction behavior. We found that AC and pStr LFP response magnitudes showed a significant positive trial-by-trial correlation for each behavioral session (Figure 7D) and the average correlation coefficients across animals did not significantly differ across sound-reward intervals (Figure 7E, p = 0.99, Kruskal-Wallis test). These results suggest the existence of strong coordination across the AC and pStr during sound processing within our task.

pStr receives direct projections from the AC, but both the pStr and AC receive direct projections from the medial geniculate body (Chen et al., 2019; Huerta-Ocampo et al., 2014; Hunnicutt et al., 2016; Smeal et al., 2008). We next sought to test to what extent this anatomical projection pattern is reflected in the temporal relationship of AC and pStr sound-evoked responses. To this end, we calculated trial-wise cross-correlations of LFP responses in AC and pStr over a 2 s sound period, from 0.5s prior to sound onset till sound termination (Figure 7F and G). The trial-averaged cross-correlation showed a clear peak at 0 lag, indicating strong synchrony in activity across the brain regions, as expected from the shared common input from the MGB (Figure 7G, black trace). However, we noticed that the shape of these average cross-correlation curves was not symmetric around 0. To test this, we compared them to cross-correlations generated after randomly shuffling the AC/pStr identity of the traces (see Methods). This form of shuffling generated cross correlations with no temporal directionality across the brain regions by design (Figure 7G, red trace). In comparison to these shuffled cross-correlations, the real cross correlations showed higher values at positive lags, indicating a temporal lead of AC relative to pStr (Figure 7G). To quantify this, we calculated the difference traces of the real and shuffled cross correlations (Figure 7H). Across the data, the median peak time of these difference traces was 38 ms and was significantly different from 0 s (Figure 7I, Wilcoxon signed rank test, p = 0.0084). Further, these peak times of difference traces were not significantly different across sound-reward intervals (Figure 7J, p = 0.94, Kruskal-Wallis test). This positive median peak time indicates that in addition to the no-lag synchrony across these brain regions, AC leads pStr during the sound predicting the time to reward in our task.

## Discussion

In this study, we established a novel behavioral paradigm based on an extension of the classical appetitive trace-conditioning task to assess sound-triggered reward time prediction in mice. Using this behavioral paradigm, we found that mice can use a sound cue to reliably predict time intervals to reward at a 1-second temporal resolution and that the ability to predict delayed reward is dependent on AC. We further found that mice use the sound onset to estimate the time to reward and that the AC LFP responses to sound onset predicted the amount of time to reward. As a downstream pathway to non-auditory brain areas, we found that the neural projections from AC to pStr, as well as the pStr itself, are necessary for sound-triggered delayed reward prediction. Sound responses in pStr also varied based on the time to reward. Finally, using simultaneous recordings in AC and pStr during performance of this task, we found strong coordination of neural activity across these brain regions, with synchronous, as well as AC-leading components of cross-correlation. Together, our findings identify AC-pStr mechanisms for sound-triggered prediction of reward timing.

Decades of research has yielded several competing models for how the brain represents time. The most explored theory underlying the timing mechanisms in animals and humans assumes the existence of an internal clock based on neural counting (Fung et al., 2021; Hinton & Meck, 1997; Leow & Grahn, 2014; Treisman, 1963; Wearden, 2005). According to this theory, higher order “centralized” clock brain regions receive inputs from various modalities and maintain a representation of time. On the other hand, parallel work in this domain supports the existence of multiple timing mechanisms distributed across different brain regions and circuits that are engaged based on the task design, sensory modality used, and the temporal resolution of the task (Bueti, 2011; Jazayeri & Shadlen, 2015; Tallot & Doyère, 2020; Matell, et al., 2011). Our findings that sound-triggered reward time prediction ability in mice at a 1-second temporal resolution is dependent on AC, strongly supports the distributed modality-specific timing model. By pharmacologically inactivating AC, we show an impairment in this ability to predict delayed reward time based on a sound cue. Further, we found that the magnitude of AC responses to the sound cue in this task encode and maintain a neural representation of the time from the sound onset to reward. These results provide evidence for an auditory-modality specific timing mechanism for sound-triggered time estimation to an imminent salient event.

Previous studies have identified different forms of neurophysiological signatures of interval timing across various brain regions (Tallot & Doyère, 2020). A group of studies in rodent visual cortex (Hussain Shuler & Bear, 2006; Namboodiri et al., 2015), basal amygdala (Pendyam et al., 2013) and basal ganglia (Hikosaka et al., 1989) have found coding of interval timing via sustained increase or decrease in spiking activity from cue onset till upcoming salient event (reward or foot shock). Other studies, primarily in the prefrontal cortex (PFC) and dorsal striatum, have found gradual firing rate ramping up as the expectation of the animal for an upcoming salient stimulus increases (Armony et al., 1998; Matell et al., 2003; Narayanan & Laubach, 2009). Another set of studies have found encoding of time intervals by phasic increases in neural activity at the time of anticipated reward in neurons in the PFC (Yumoto et al., 2011), dopaminergic neurons in the ventral tegmental area (Fiorillo et al., 2008) and in the visual cortex (Hussain Shuler & Bear, 2006). A common factor across most of these studies is that the amount of time to the salient event is encoded separately from the sensory stimulus that triggers its anticipation. In contrast, we find that the magnitude of the AC responses to the sound itself also encode the anticipated time to reward. Thus, the cue and the reward-timing prediction associated with it are jointly coded in the AC. This form of integration between sound and the temporal expectation associated with it is consistent with a previous study, in which the AC responses to a tone were found to be different when rats expected a target sound to be presented either immediately after the tone or about 1 second later (Jaramillo & Zador, 2011). Our findings extend and generalize these results to demonstrate the existence of timing prediction of non-auditory cues within the AC, and that interval timing duration is represented in a continuous manner within the scale of 0.5-5s.

To investigate how the auditory corticostriatal projections are involved in sound-triggered time estimation behavior, we used chemogenetic inactivation to establish the necessity of the anterograde projections from AC to pStr in this task. We acquired simultaneous LFP recordings from AC and pStr to show that neural activity in these regions is temporally synchronized during the sound cue predicting the interval timing to reward. The activity in AC and pStr was highly synchronized at 0s lag, which is consistent with the common input received by both these regions from the medial geniculate body (MGB) (Chen et al., 2019; Huerta-Ocampo et al., 2014; Hunnicutt et al., 2016; Smeal et al., 2008). In the context of our task, this would suggest that the sound-triggered reward timing encoded in AC and in pStr could be simultaneously and differentially modulated by the neural inputs from MGB. In addition to the direct projections from MGB, pStr receives strong monosynaptic projections from AC (Znamenskiy and Zador, 2013; Xiong et al., 2015; Huang et al., 2023). This is consistent with our finding that in addition to the synchronous activity across these brain regions, a second component of the cross-correlograms points at AC leading the pStr by 38ms on average. Our LFP recordings in AC and pStr provided a quantification of the overall functional and temporal communication patterns across the AC and pStr. Future studies using single unit recordings with cell-type specificity could elucidate how these phenomena are represented at the local ensemble level.

While sound-triggered action timing on the scale of seconds is important for many everyday behaviors, it is a particularly critical ability in humans for verbal communication. In verbal communication, humans use incoming speech sounds to predict when and what sounds are expected to follow (Heinks-Maldonado et al., 2005, 2006; Nixon & Tomaschek, 2021; Tremblay et al., 2013). Furthermore, the sequential nature of speech sounds helps arrange incoming information in a temporal structure and this ability is impaired when there are hearing deficits (Füllgrabe, 2013; Grose & Mamo, 2010; Helfer & Jesse, 2021; Ozmeral et al., 2016; Tallal et al., 1995). Deaf or hard of hearing children face challenges in sequential time perception, which impairs their storytelling ability, a key cognitive development factor (Eden & Leibovitz-Ganon, 2022). In comparison, children with postlingual cochlear implants exhibit greater improvements in time perception and consequently, their storytelling ability, emphasizing the role of hearing acquisition in sound-guided temporal processing. Other research involving children with mild hearing loss have showcased the benefits of interventions on time sequencing and storytelling (Ingber & Eden, 2011). These studies collectively underscore the critical role of sound-guided timekeeping in developing and maintaining language-related skills and highlight the need to further study the neural mechanisms underlying these processes to develop tailored support for individuals with hearing impairments.

## Materials and Methods

All animal procedures were performed in accordance with the University of Michigan animal care committee’s regulations.

### Animals

We used 56 (38 males, 18 females, 8-16 weeks of age) C57BL/6J mice (Jax number: 000664). Mice were individually housed under a reverse 12h light/12h dark cycle, with lights on at 8:30 pm and off at 8:30 am, and had access to ad libitum food, water, and enrichment. During the behavioral training period, mice were on water restriction and given 2-5ml of water each day.

### Surgical procedure

All surgeries were performed on mice anesthetized using isoflurane (1.5-2% vol/vol). Anesthetized mice were placed in a stereotaxic frame (Kopf 514 Instruments, CA, USA), and an anti-inflammatory drug (Carprofen, 5mg/kg, subcutaneous injection) and a local anesthetic (lidocaine, subcutaneous injection) were administered. A custom-made lightweight (<1 gr) titanium head bar was attached to the back of the skull using dental cement and cyanoacrylate glue to allow for head-fixed behavior. During the surgery, body temperature was maintained at 38^ᵒ^C, and the depth of anesthesia was regularly assessed by checking the pinch withdrawal reflex. A small craniotomy was performed over target coordinates relative to the bregma (AC: −2.7mm anterior, ±4.3mm from midline, −0.55mm ventral, 0° angle; pStr: −1.7mm anterior, ±3.35mm from midline, −2.8mm ventral, 0° angle).

To chemogenetically target the neural projections from AC to pStr and for pharmacological inactivation experiments using muscimol, custom-made cannulae (25-gauge tubing) or guide cannulas (Plastics One) were placed at the surface of the brain in these craniotomies at the target regions and secured to the skull using dental cement. Dummy cannulae were inserted into these cannulae to prevent outside debris from entering the cannula.

Mice were treated with Carprofen for 48 hours post-surgically and were allowed to recover for a week.

### Electrophysiological recordings

Tungsten wire electrodes (two 50 μm wires bundle) with < 50kΩ of impedance were used to acquire local field potential (LFP) responses in AC and pStr. Electrodes were dipped in DiI dye (Invitrogen, Catalog # 22885) before insertion. The ground screw was positioned over the cerebellum (2.0 mm posterior to lambda, 3.5 mm from midline). Electrodes were unilaterally implanted into AC and pStr on the right hemisphere, (distance between the two electrodes within a region was <50 μm, distance between electrode arrays in AC and pStr was ∼2.67 mm). LFP signals were acquired using a Tucker-Davis Technologies (TDT) acquisition system and Synapse Lite Software. The output bioelectrical signal was digitized, sampled at 6 kHz, and bandpass filtered in 0.5-300Hz for LFP recordings. All data acquired was saved for offline data processing.

### Virus injections

Viruses were acquired from Addgene to inactivate the anterograde projections from AC to pStr chemogenetically. To achieve projection specificity in C57BL/6J WT mice, we used an established dual viral approach as shown in Figure 5A. We bilaterally injected Cre-dependent DREADD virus (Roth, 2016): AAV5-hSyn-DIO-hM4D(Gi)-mCherry (2.4E+13 vg/ml, 350nl, Addgene catalog # 44362) or AAV5-hSyn-DIO-mCherry (2.6E+13 vg/ml, 350nl, Addgene catalog # 50459) into the AC and a retrograde Cre virus: pENN/AAVrg-hSyn-Cre-WPRE-hGH (1.8E+13 vg/ml, 200nl, Addgene catalog #105553) into the pStr. The infusions were done using a 32-gauge injection needle (customed-length per infusion site) or through thin internal cannulas (Plastics One) inserted into the implanted cannulae in the brain, connected to a 10 μl Hamilton syringe at a rate of 50nl/min.

### Drug administration

#### Muscimol infusions

Mildly sedated mice were bilaterally infused with 0.5µg/µl muscimol (BODIPY TMR-X fluorophore-conjugated, ThermoFisher, Catalog Number – M23400) dissolved in phosphate-buffered saline (PBS) and 1.5% DMSO or PBS with 1.5% DMSO as a control (Volume per hemisphere = AC: 750nl (Aizenberg et al., 2015; Sun et al., 2022), pStr: 360nl (Guo et al., 2018)) at a rate of 150nl/min, into cannulae implanted in target sites. The infusions were done via custom-made injectors or thin internal cannulas (Plastics One) as previously described.

#### CNO injections

5mg Clozapine-N-Oxide (CNO) (HelloBio) was diluted in 0.9% saline solution. All animals in the chemogenetic inactivation experiments were first injected with saline (5mg/kg, i.p.) as a control and then were injected with the prepared CNO (5mg/kg, i.p.) solution the following day, to chemogenetically inactivate the projections from AC to pStr.

### Behavior

All our behavioral setups were custom built and controlled by an Arduino (Arduino Uno board with an Adafruit Music Maker shield) circuit. Behavioral data acquired through the Arduino IDE software was saved in text files for analysis. Videos of animal behavior were acquired using Logitech C920 HD Pro camera on the LogiCapture software and using the Angetube 1080p web camera on the Bandicam software, under red light conditions.

#### Paradigm

Mice were trained on an appetitive sound-triggered reward time prediction task. In this task, mice on water restriction were head-fixed inside a tube to reduce movement-related artifacts, presented a sound cue from a speaker (4Ω, 3W magnetic speakers, placed ∼10cm away from the animal’s head on the left) and trained to consume a water reward from a reward port placed close to its mouth. A trial constituted a 1.5s long sound cue and a water reward delivery separated by a fixed time interval (0.5-5s), with randomized inter-trial intervals in the range of 2-6s (Figure 1A, Trial block). The sound cue was a sequence of three 0.5s long pure tones (8kHz, 12kHz, 16kHz; 5ms rise/fall time) generated at a 25kHz sampling rate using MATLAB (Mathworks 2019a).

#### Training

Water-restricted mice were handled and habituated to the experimental setup for ∼7 days. In this period, mice were head fixed and trained to lick the reward port through which a water reward was delivered randomly at 3-10s intervals without any sound cue. The reward port consisted of a metal tube that delivered a fixed amount of water (∼3ul) each trial, connected to a capacitance-based lick detector that allowed recording individual lick times. Mice were also familiarized with the sound cue used in behavioral training over the last 3 days of habituation through random sound presentations (∼50 times across all 3 days) at 3-10s intervals without any reward delivery.

Habituated mice started training on trials with a fixed time interval of 1.5s from sound termination time to reward, with 150-250 trials per daily training session. After 7-10 days of training, catch trials in which reward was withheld were randomly introduced 15-20% of the trials/session. Once the animal learnt to predict reward time at 1.5s interval, it was then trained to use the same sound to predict a different interval of time between sound and reward (sound-reward interval). We trained each animal to predict timed reward using the same sound cue at four sound-reward intervals – 0.5s, 1.5s, 2.5s and 5s and always trained them to learn the time intervals in this order – 1.5s ◊ 2.5s ◊ 5s ◊ 0.5s (Figure 1A).

To identify whether mice predicted the sound-reward interval duration from sound onset or sound termination, we tested a subset of mice trained to predict reward at 1.5s from sound cue, to use a shorter duration sound cue to predict reward. On this testing day, we randomly interspersed 35% of the trials with 1s long sound cue (a sequence of 8kHz and 12kHz pure tones, each 0.5s long, with 5ms rise/fall time) to deliver reward at the same time interval – short sound trials (Figure 3A), along with standard 1.5s long sound cue trials.

#### Behavioral training for the pharmacological inactivation experiments

In these experiments, we used a GABA-A receptor agonist, muscimol, to inactivate AC or pStr to establish their causal role in sound-triggered reward time prediction task. Each of these experiments consisted of two different cohorts of mice. One cohort of animals were trained to predict timed reward at 1.5s from sound termination (Figures 2, 5, and 6; 1.5s Delay task) and another cohort of animals were trained on an alternative version of the task where reward immediately followed the sound cue (Figures 2, 5, and 6; No-Delay task).

#### Behavioral training for acquiring electrophysiological recordings in AC and pStr

Mice implanted with wire electrodes underwent the same habituation protocol as described above. In this habituation phase, these mice were habituated to cables plugged to the electrode connectors on their head implants and trained to lick the reward port to consume the water reward. The reward port for these experiments was a tube fitted with an IR sensor to detect licks through beam breaks. Following habituation, they were trained on the previously described sound-triggered reward time prediction task on the 4 different time intervals between sound and reward in the order 1.5s ◊ 2.5s ◊ 5s ◊ 0.5s, while their LFP responses in AC and pStr were recorded throughout each daily training session. LFP responses were monitored for movement using video recording and periods of movement were eliminated prior to analysis.

#### Behavioral training for chemogenetic inactivation of AC-pStr projections experiments

We used chemogenetic inactivation of anterograde projections from AC to pStr to identify their role in sound-triggered reward time prediction task. Animals in these experiments started by learning to predict timed reward at 1.5s from sound (1.5s Delay task) and then underwent chemogenetic AC-pStr projection inactivation with CNO injection to test for effect on behavioral performance. After a 4-day washout period, a subset of these same animals underwent training to predict sound-guided timed reward at 5s interval (5s Delay task) and chemogenetic inactivation with CNO injection to test for behavioral effect to predict timed reward at 5s from sound. Following another 4-day washout period, these animals trained and underwent chemogenetic manipulation on the No-Delay task (Figure 5A, Task timeline).

In a subset of animals in which DREADDs were expressed in the AC-pStr projections, we also simultaneously recorded LFP responses in pStr using tungsten wire electrodes, while they trained on the 1.5s Delay and No-Delay tasks.

### Data analysis

All analyses were done using custom-written MATLAB (Mathworks 2022a) scripts unless otherwise mentioned.

#### Behavioral data analysis

To quantify the animal’s ability to predict reward at a fixed time interval from a sound cue, we extracted individual lick times per trial and averaged these licks across trials for each daily training session to get a predictive licking response curve. To measure learning, we computed the slope of this predictive licking response curve (MATLAB command: polyfit, degree 1) in the predictive lick period, which was defined differently for rewarded and catch trials. For rewarded trials, the predictive lick period was defined as the time from sound termination to 100ms prior to reward delivery time for each sound-reward interval. Contrastingly, we used a more conservative definition of the predictive lick period for catch trials, using the period from 200ms prior to sound termination to the time at which reward was expected for the 0.5s interval, for all the sound-reward intervals. We used this slope measure in catch trials to ascertain when the animals had learnt to consistently predict reward within each sound-reward time interval. We compared the slopes of the predictive licking curve for catch trials across training days for each sound-reward interval and picked those days for which the slope value crossed the slope threshold (mean + 3*standard deviation of slope values across all training days per sound-reward interval). Amongst the training days that satisfied this criterion, the day with maximum slope value was chosen as the “best” behavior day for each sound-reward interval per animal. Additionally, we computed the full width at half maxima of the predictive licking response curve over the predictive lick period defined above for catch trials to estimate the precision of the animal’s ability to predictive lick for each sound-reward interval (Supplementary Figure 1).

We determined the effect of muscimol and chemogenetic inactivation on behavioral performance by comparing the predictive lick responses in catch trials on control training day (PBS infusion for pharmacological inactivation experiments or saline injections (i.p.) for chemogenetic inactivation experiments) to the predictive lick responses on the manipulation day (muscimol (MUS) infusion for pharmacological inactivation experiments or CNO injections (i.p.) for chemogenetic inactivation experiments) using this formula –

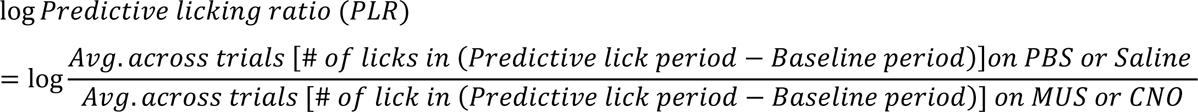

*Where, baseline period = sound onset time – 750ms to sound onset time, and predictive lick period for delay tasks = reward time – 250ms to reward time + 500ms for no-delay task = sound onset time + 750ms*.

To check whether animals’ ability to lick for reward changed based on sound termination time or not, we compared their predictive lick responses in standard and short sound trials using a variation of the PLR described above –

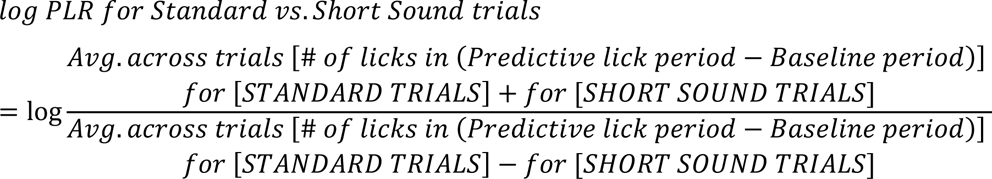

*Where, baseline period = sound onset time – 750ms to sound onset time, and predictive lick period = sound offset time to reward time-100s*

#### Electrophysiological data processing and analysis

Acquired electrophysiological data was extracted using TDTBin2mat script (provided by TDT) and organized to synchronize it with lick response times for each session across sound-reward intervals per animal. Using slopes of the predictive lick response curve of catch trials, the “best” behavior day was determined for each sound-reward interval per animal, as described above. Trials in each training session with no licks in the 200ms period from sound onset or in the 500ms period from sound offset (hereafter referred to as no-lick trials) were extracted and further analysis was carried out only on these no-lick trials on the “best” behavior days for each sound-reward interval across animals. Session-wise LFP signals for AC and pStr were filtered for movement-related artifacts by eliminating any signal above a threshold of mean LFP signal for the session + 3*standard deviation of the session LFP signal and then were z-scored for each session prior to analysis.

To analyze sound-evoked LFP responses, trial-wise LFP activity in AC and pStr were aligned to the sound onset and baseline-corrected by subtracting its average during the 10ms period from sound onset for computing the onset response magnitude (similar to (Taaseh et al., 2011)) or from sound offset for computing the offset response magnitude. Response magnitude was defined as the amplitude of the maximum trough (most negative) in the 40ms period from sound onset/offset for each no-lick trial per session. Response magnitudes were averaged across trials per session and normalized to the average response magnitude of the shortest sound-reward time interval (0.5s) session per animal. Normalized response magnitudes were combined across animals per sound-reward interval and compared using the Kruskal-Wallis test (MATLAB command: kruskalwallis), with a post-hoc Tukey-Kramer test to determine individual group differences (MATLAB command: multcompare applied on the kruskalwallis output).

To compare the pStr onset response magnitudes in saline and CNO conditions, we averaged the response magnitudes across trials and tested for significant differences per animal between the conditions using the Wilcoxon rank-sum test (MATLAB command: ranksum). These average response magnitudes were then normalized to average of the saline condition to determine the population-level trends for the 1.5 Delay and No-Delay tasks.

We examined the coordination in AC and pStr LFP activity during sound by computing the trial-by-trial correlation of sound onset response magnitudes in AC and pStr (MATLAB command: corrcoef). Significant correlation coefficients across animals were combined per sound-reward interval and compared across intervals using the Kruskal-Wallis test (MATLAB command: kruskalwallis), with a post-hoc Tukey-Kramer test to determine individual group differences (MATLAB command: multcompare applied on the kruskalwallis output).

To determine the temporal relationship of AC and pStr LFP responses during sound-triggered reward time prediction behavior, we ran a cross-correlation analysis of the LFP responses in AC and pStr for each session (MATLAB command: xcorr, using the “normalized” model, maxlag = ±100ms) over a 2s period from 0.5s prior to sound onset to sound termination time. By shuffling the identity of AC and pStr traces per session and computing their cross-correlation coefficients, we simulated chance cross correlation coefficients between AC and pStr sound responses (Number of iterations = 100). We generated difference traces by calculating the difference in average cross correlation coefficients between real and simulated data, and then determined the time which these difference traces peaked giving us the time lag between AC and pStr sound responses per training session. We calculated the median of this peak time of difference traces across all animals and sound-reward intervals and compared this median against 0s using the Wilcoxon signed rank test (MATLAB command: signrank).

#### Statistical tests

We used statistical tests at a p<0.05 significance level and α = 0.05 for all comparisons unless otherwise indicated.

### Histology

Mice were euthanized with an overdose of isoflurane (5%) or carbon dioxide (2%) and perfused transcardially with PBS (0.9%), followed by 10% paraformaldehyde (PFA). The brain tissue was removed and fixed in 10% PFA for 72 hours. For cryoprotection, the brain tissue was transferred to 30% sucrose solution for 3-4 days before sectioning. Coronal sections (50µm thickness) were obtained in a cryostat (Leica) and kept in PBS at 4°C before mounting. All sections were mounted in glass slides and covered with Fluoroshield mounting medium with DAPI (Abcam, USA). Images of brain sections were acquired using a fluorescent microscope (Zeiss) equipped with an apotome (Olympus) and the ZenPro software.

For all histological validations, brain sections were imaged with a 10x objective, examined for cell nuclei labeling DAPI expression (470 nm), and saved as both multichannel and individual fluorophore channels composite tiff images. Histological validation was done by overlaying brain sections over slice images from the Allen Institute’s 10 μm voxel 2017 version from the Allen Mouse Brain Common Coordinate Framework (Wang et al., 2020). Histological validation was used as an inclusion criterion for all behavioral, electrophysiological, and chemogenetic experiments.

#### Tungsten electrode tracks verification

Electrode tracks were identified by DiI expression (550 nm). Images with DiI expression and electrode tracks were saved and further analyzed with the open-source SHARP-Track toolkit (Shamash et al., 2018). Verified electrode tip coordinates were compared to the coordinate range showing the projections from AC to pStr shown in the Allen Mouse Brain Connectivity Atlas (Allen Mouse Brain Connectivity Atlas, connectivity.brain-map.org/projection/experiment/146858006).

#### Reconstruction of virus expression

The procedure for reconstructing viral expression volumes was similar to electrode track reconstruction. However, the regions defined in each slice are two-dimensional, and the volume is delineated at the 3D projection of all the combined sections showing virus expression. AP, ML, and DV coordinates from all sections with positive mCherry expression were again compared to the range of AC-pStr projections shown in the Allen Mouse Brain Connectivity Atlas (Allen Mouse Brain Connectivity Atlas, connectivity.brain-map.org/projection/experiment/146858006). Animals showing mCherry cell body expression outside AC areas and mCherry axonal expression outside pStr were not included.

#### Muscimol infusion

To verify the muscimol diffusion within the target areas, brain slices were examined for BODYPY-TmX (BDP-T, 573 nm) expression. Brain sections showing positive BDP-T labeling were saved per animal. Further comparison with the reference Allen Atlas and 3D projection showed the AP and ML coordinates with muscimol diffusion. Clear hit into AC or pStr was considered a diffusion only within the target areas. For pStr experiments, animals with muscimol diffusion up to rostral striatal areas (anterior to −1.2mm from bregma) were not included.

## Acknowledgements

We would like to thank Lily Allen, Zeinab Mansi, Lindsay Cain, and Grant Griesman for their help with training animals on the task and Siddharth Mohite for his help with setting up the analysis pipeline scripts.

This work was supported by a National Institute of Health grant R01NS129874 (G.R.), a National Institute of Health grant R01MH063649 (G.R. Co-I) and an Alzheimer’s Association Research Grant 21-850571 (G.R.)

## Author contributions

G.R. and H.S. conceived the project and designed the experiments, H.S. and K.S.P. performed surgeries, conducted experiments and histological analysis, N.A, K.G, K.L, N.N, A.O, and B.W, trained animals on the behavioral task and conducted parts of the histological analysis, H.S. performed all data analysis, G.R., and H.S. wrote the manuscript, and B.W. made the illustrations.

## Supplementary Figures

**SUPPLEMENTARY FIGURE 1.**
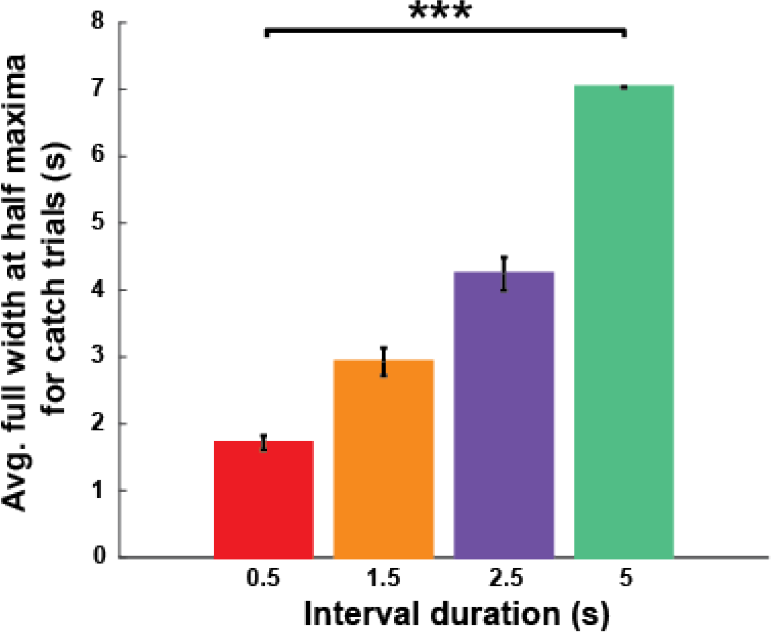
Precision of predictive licking ability across sound-reward intervals. The average full-width at half maxima of the predictive licking curves across animals was significantly different across sound-reward intervals (***p = 3.85×10^-6^, Kruskal-Wallis test).

**SUPPLEMENTARY FIGURE 2.**
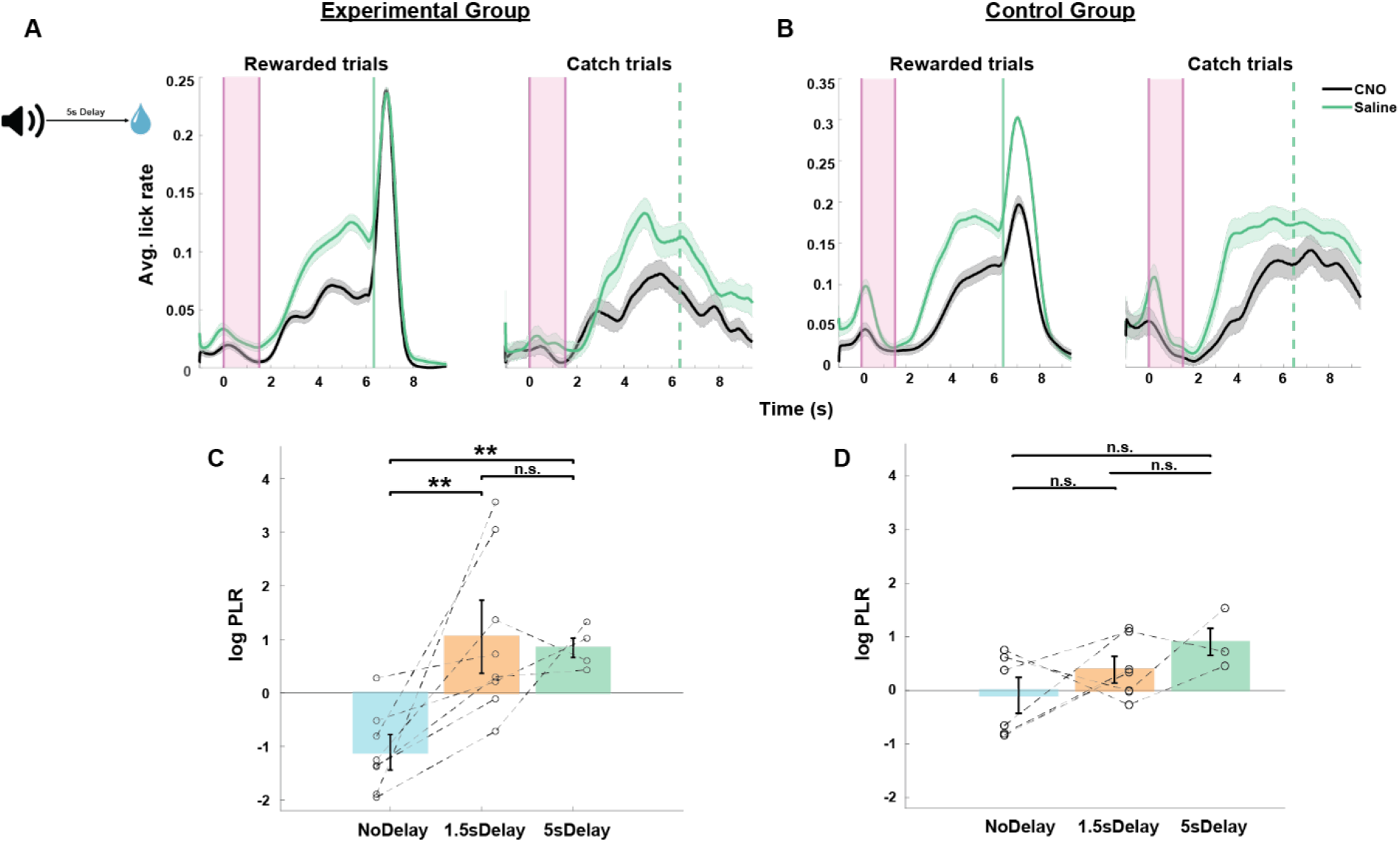
Effect of chemogenetic inactivation of AC to pStr projections on the 5s Delay task. A. and B. Average peri-sound lick rate response curves (solid line denotes mean, shaded area represents SEM across trials) of an example animal trained to predict reward at 5s sound-reward interval when injected with saline (green) and CNO (black) for rewarded trials (left) and catch trials (right). Shaded pink region represents the 1.5s long sound period. Solid and dotted green lines represent when reward was given in rewarded trials and expected in catch trials. Left column represents animal from the experimental group and right column represents animals from the control group. A. and D. Average log Predictive Licking Ratio (log PLR) across animals trained on the 1.5s Delay, 5s Delay and No-Delay tasks in the experimental group and in the control group reiterated from Fig. 5E and H. Lines connecting the circles represent the PLR for each animal when trained on the 1.5s Delay, 5s Delay and No-Delay tasks. The average PLR for experimental group animals was significantly higher for 5s Delay task than No-Delay task (**p = 0.004, Wilcoxon rank sum test) and was not significantly different for the control group animals (p = 0.167, Wilcoxon rank sum test).

**SUPPLEMENTARY FIGURE 3.**
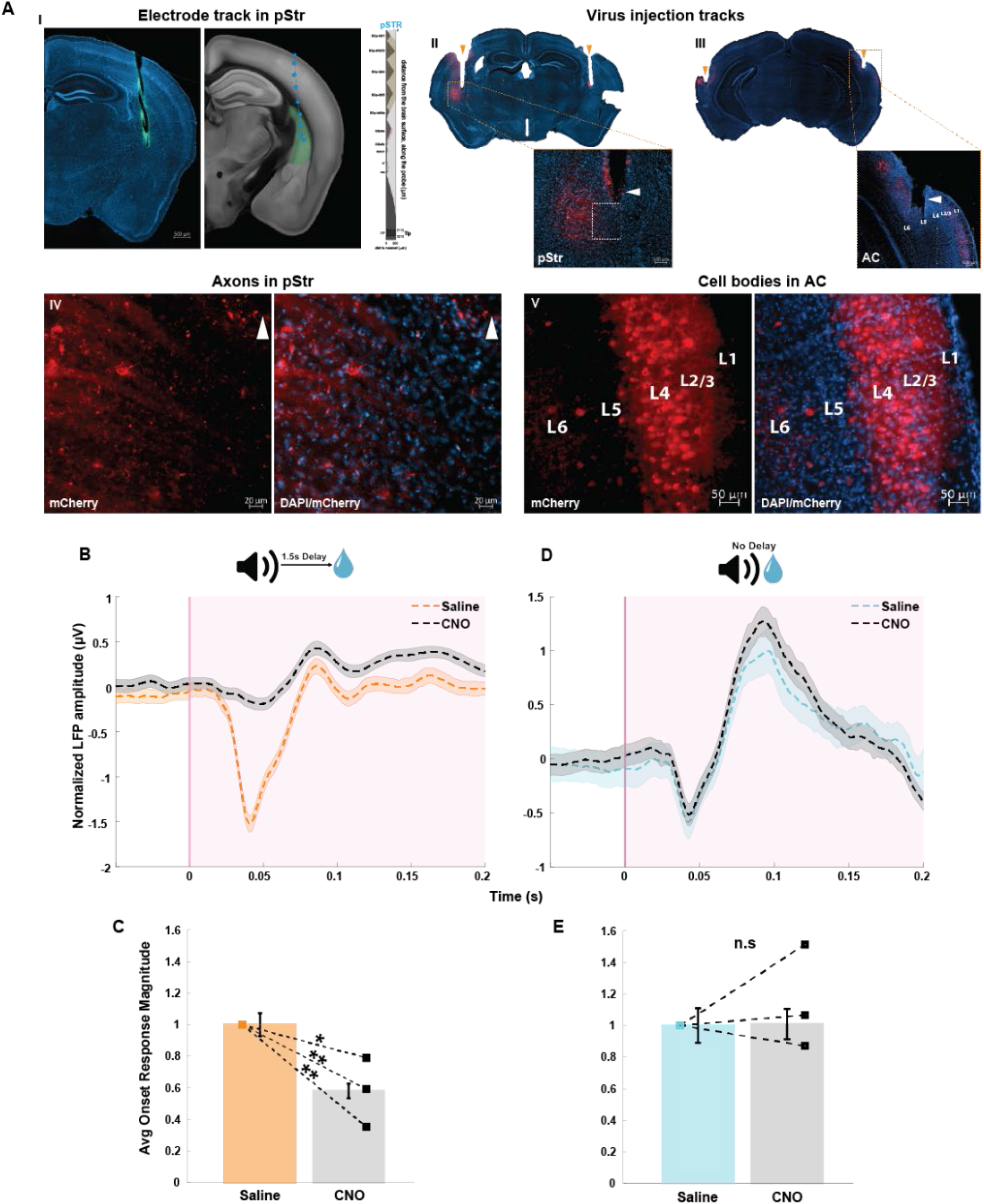
Histological verification of targeting AC to pStr projections and change in LFP responses in pStr following chemogenetic inactivation of these projections. A. Histological validation of electrode position in pStr and chemogenetic virus expression in AC to pStr projections. I. Electrode position in pStr is denoted by the green dotted lines. II. and III. Representation of the virus injection tracks in AC and pStr. IV. and V. are magnified images from II and III showing axons in pStr and cell bodies in AC. B. and D. Normalized average pStr LFP (solid line denotes mean, shaded area represents SEM across no-lick trials) recorded in response to the sound onset from an example animal trained on the 1.5s Delay task (left) and on the No-Delay task (right) with saline (orange/light blue) and CNO (black) injections. The shaded pink region represents the period from sound onset. C. and E. Comparison of the normalized average pStr onset response magnitude computed across animals (N = 3) for the 1.5s Delay task (Fig C) and for the No-Delay task (Fig E) on saline and CNO conditions. Error bars represent mean ± SEM across animals. Comparison between saline and CNO conditions yields *p<0.05 for each animal when trained on the 1.5s Delay task and was not significantly different when trained on the No-Delay task.

## References

Aizenberg, M., Mwilambwe-Tshilobo, L., Briguglio, J. J., Natan, R. G., & Geffen, M. N. (2015). Bidirectional Regulation of Innate and Learned Behaviors That Rely on Frequency Discrimination by Cortical Inhibitory Neurons. PLoS Biology, 13(12), 1–32. 10.1371/journal.pbio.1002308

Armony, J. L., Quirk, G. J., & Ledoux, J. E. (1998). Differential effects of amygdala lesions on early and late plastic components of auditory cortex spike trains during fear conditioning. Journal of Neuroscience, 18(7), 2592–2601. 10.1523/jneurosci.18-07-02592.1998

Bathellier, B., Ushakova, L., & Rumpel, S. (2012). Discrete Neocortical Dynamics Predict Behavioral Categorization of Sounds. Neuron, 76(2), 435–449. 10.1016/j.neuron.2012.07.008

Benichov, J. I., Globerson, E., & Tchernichovski, O. (2016). Finding the beat: From socially coordinated vocalizations in songbirds to rhythmic entrainment in humans. Frontiers in Human Neuroscience, 10(June), 1–7. 10.3389/fnhum.2016.00255

Benichov, J. I., & Vallentin, D. (2020). Inhibition within a premotor circuit controls the timing of vocal turn-taking in zebra finches. Nature Communications, 11(1), 1–10. 10.1038/s41467-019-13938-0

Bernal, B., & Ardila, A. (2016). From hearing sounds to recognizing phonemes: Primary auditory cortex is a truly perceptual language area. AIMS Neuroscience, 3(4), 454–473. 10.3934/Neuroscience.2016.4.454

Bertero, A., Zurita, H., Normandin, M., & Apicella, A. J. (2020). Auditory Long-Range Parvalbumin Cortico-Striatal Neurons. Frontiers in Neural Circuits, 14(July), 1–15. 10.3389/fncir.2020.00045

Bueti, D. (2011). The sensory representation of time. Frontiers in Integrative Neuroscience, 5(August), 1–3. 10.3389/fnint.2011.00034

Bueti, D., Bahrami, B., & Walsh, V. (2008). Sensory and association cortex in time perception. Journal of Cognitive Neuroscience, 20(6), 1054–1062. 10.1162/jocn.2008.20060

Buhusi, C. V., & Meck, W. H. (2005). What makes us tick? Functional and neural mechanisms of interval timing. Nature Reviews Neuroscience, 6(10), 755–765. 10.1038/nrn1764

Chen, L., Wang, X., Ge, S., & Xiong, Q. (2019). Medial geniculate body and primary auditory cortex differentially contribute to striatal sound representations. Nature Communications, 10(1), 1–10. 10.1038/s41467-019-08350-7

Chubykin, A. A., Roach, E. B., Bear, M. F., & Shuler, M. G. H. (2013). A Cholinergic Mechanism for Reward Timing within Primary Visual Cortex. Neuron, 77(4), 723– 735. 10.1016/j.neuron.2012.12.039

Cook, J. R., Li, H., Nguyen, B., Huang, H. H., Mahdavian, P., Kirchgessner, M. A., Strassmann, P., Engelhardt, M., Callaway, E. M., & Jin, X. (2022). Secondary auditory cortex mediates a sensorimotor mechanism for action timing. Nature Neuroscience, 25(3), 330–344. 10.1038/s41593-022-01025-5

Dunlap, A. G., Besosa, C., Pascual, L. M., Chong, K. K., Walum, H., Kacsoh, D. B., Tankeu, B. B., Lu, K., & Liu, R. C. (2020). Becoming a better parent: Mice learn sounds that improve a stereotyped maternal behavior. Hormones and Behavior, 124(July), 104779. 10.1016/j.yhbeh.2020.104779

Eden, S., & Leibovitz-Ganon, K. (2022). The effects of cochlear implants on sequential time perception. Deafness and Education International, 24(2), 160–178. 10.1080/14643154.2021.1902644

Ehret, G. (2005). Infant rodent ultrasounds - A gate to the understanding of sound communication. Behavior Genetics, 35(1), 19–29. 10.1007/s10519-004-0853-8

Fiorillo, C. D., Newsome, W. T., & Schultz, W. (2008). The temporal precision of reward prediction in dopamine neurons. Nature Neuroscience, 11(8), 966–973. 10.1038/nn.2159

Francis, N. A., Winkowski, D. E., Sheikhattar, A., Armengol, K., Babadi, B., & Kanold, P. O. (2018). Small Networks Encode Decision-Making in Primary Auditory Cortex. Neuron, 97(4), 885–897.e6. 10.1016/j.neuron.2018.01.019

Fritz, J., Elhilali, M., & Shamma, S. (2005). Active listening: Task-dependent plasticity of spectrotemporal receptive fields in primary auditory cortex. Hearing Research, 206(1–2), 159–176. 10.1016/j.heares.2005.01.015

Fritz, J., Shamma, S., Elhilali, M., & Klein, D. (2003). Rapid task-related plasticity of spectrotemporal receptive fields in primary auditory cortex. Nature Neuroscience, 6(11), 1216–1223. 10.1038/nn1141

Füllgrabe, C. (2013). Age-dependent changes in temporal-fine-structure processing in the absence of peripheral hearing loss. American Journal of Audiology, 22(2), 313–315. 10.1044/1059-0889(2013/12-0070)

Fung, B. J., Sutlief, E., & Hussain Shuler, M. G. (2021). Dopamine and the interdependency of time perception and reward. Neuroscience and Biobehavioral Reviews, 125, 380–391. 10.1016/j.neubiorev.2021.02.030

Geissler, D. B., & Ehret, G. (2004). Auditory perception vs. recognition: Representation of complex communication sounds in the mouse auditory cortical fields. European Journal of Neuroscience, 19(4), 1027–1040. 10.1111/j.1460-9568.2004.03205.x

Grose, J. H., & Mamo, S. K. (2010). Processing of temporal fine structure as a function of age. Ear and Hearing, 31(6), 755–760. 10.1097/AUD.0b013e3181e627e7

Guo, L., Walker, W. I., Ponvert, N. D., Penix, P. L., & Jaramillo, S. (2018). Stable representation of sounds in the posterior striatum during flexible auditory decisions. Nature Communications, 9(1). 10.1038/s41467-018-03994-3

Guo, L., Weems, J. T., Walker, W. I., Levichev, A., & Jaramillo, S. (2019). Choice-selective neurons in the auditory cortex and in its striatal target encode reward expectation. Journal of Neuroscience, 39(19), 3687–3697. 10.1523/JNEUROSCI.2585-18.2019

Heilbron, M., & Chait, M. (2018). Great Expectations: Is there Evidence for Predictive Coding in Auditory Cortex? Neuroscience, 389, 54–73. 10.1016/j.neuroscience.2017.07.061

Heinks-Maldonado, T. H., Mathalon, D. H., Gray, M., & Ford, J. M. (2005). Fine-tuning of auditory cortex during speech production. Psychophysiology, 42(2), 180–190. 10.1111/j.1469-8986.2005.00272.x

Heinks-Maldonado, T. H., Nagarajan, S. S., & Houde, J. F. (2006). Magnetoencephalographic evidence for a precise forward model in speech production. NeuroReport, 17(13), 1375–1379. 10.1097/01.wnr.0000233102.43526.e9

Helfer, K. S., & Jesse, A. (2021). Hearing and speech processing in midlife. Hearing Research, 402, 108097. 10.1016/j.heares.2020.108097

Hikosaka, O., Sakamoto, M., & Usui, S. (1989). Functional properties of monkey caudate neurons. III. Activities related to expectation of target and reward. Journal of Neurophysiology, 61(4), 814–832. 10.1152/jn.1989.61.4.814

Hinton, S. C., & Meck, W. H. (1997). Erratum: The “internal clocks” of circadian and interval timing. Endeavour, 21(2), 82–87. 10.1016/S0160-9327(97)01043-0

Huang, W., Wang, Y., Qin, J., He, C., Li, Y., Wang, Y., Li, M., Lyu, J., Zhou, Z., Jia, H., Pakan, J., Xie, P., & Zhang, J. (2023). A corticostriatal projection for sound-evoked and anticipatory motor behavior following temporal expectation. NeuroReport, 34(1), 1–8. 10.1097/WNR.0000000000001851

Huerta-Ocampo, I., Mena-Segovia, J., & Bolam, J. P. (2014). Convergence of cortical and thalamic input to direct and indirect pathway medium spiny neurons in the striatum. Brain Structure & Function, 219(5), 1787–1800. 10.1007/s00429-013-0601-z

Hunnicutt, B. J., Jongbloets, B. C., Birdsong, W. T., Gertz, K. J., Zhong, H., & Mao, T. (2016). A comprehensive excitatory input map of the striatum reveals novel functional organization. ELife, 5(November2016), 1–32. 10.7554/eLife.19103

Hussain Shuler, M. G., & Bear, M. F. (2006). Reward Timing in the Primary Visual Cortex. Scientific Reports, March, 231–237. 10.4324/9780203181232-18

Ingber, S., & Eden, S. (2011). Enhancing Sequential Time Perception and Storytelling Ability of Deaf and Hard of Hearing Children. American Annals of the Deaf, 156, 391–401.

Jaramillo, S., & Zador, A. M. (2011). The auditory cortex mediates the perceptual effects of acoustic temporal expectation. Nature Neuroscience, 14(2), 246–253. 10.1038/nn.2688

Jasmin, K., Lima, C. F., & Scott, S. K. (2019). Understanding rostral–caudal auditory cortex contributions to auditory perception. Nature Reviews Neuroscience, 20(7), 425–434. 10.1038/s41583-019-0160-2

Jazayeri, M., & Shadlen, M. N. (2015). A Neural Mechanism for Sensing and Reproducing a Time Interval. Current Biology, 25(20), 2599–2609. 10.1016/j.cub.2015.08.038

Jones, C. R. G., & Jahanshahi, M. (2011). Dopamine modulates striato-frontal functioning during temporal processing. Frontiers in Integrative Neuroscience, 5(70).

Khouri, L., & Nelken, I. (2015). Detecting the unexpected. Current Opinion in Neurobiology, 35, 142–147. 10.1016/j.conb.2015.08.003

King, A. J., & Schnupp, J. W. H. (2007). The auditory cortex. Current Biology, 17(7), 236–239. 10.1016/j.cub.2007.01.046

King, A. J., Teki, S., & Willmore, B. D. B. (2018). Recent advances in understanding the auditory cortex. F1000Research, 7(1555). 10.12688/F1000RESEARCH.15580.1

Kuchibhotla, K., & Bathellier, B. (2018). Neural encoding of sensory and behavioral complexity in the auditory cortex. Current Opinion in Neurobiology, 52, 65–71. 10.1016/j.conb.2018.04.002

Kuchibhotla, K. V., Gill, J. V., Lindsay, G. W., Papadoyannis, E. S., Field, R. E., Sten, T. A. H., Miller, K. D., & Froemke, R. C. (2017). Parallel processing by cortical inhibition enables context-dependent behavior. Nature Neuroscience, 20(1), 62–71. 10.1038/nn.4436

Kurti, A. N., & Matell, M. S. (2011). Nucleus Accumbens Dopamine Modulates Response Rate but Not Response Timing in an Interval Timing Task. Behavioral Neuroscience, 125(2), 215–225. 10.1037/a0022892

LeDoux, J. E., Farb, C. R., & Romanski, L. M. (1991). Overlapping projections to the amygdala and striatum from auditory processing areas of the thalamus and cortex. Neuroscience Letters, 134(1), 139–144. 10.1016/0304-3940(91)90526-Y

Lee, J., & Rothschild, G. (2021). Encoding of acquired sound-sequence salience by auditory cortical offset responses. Cell Reports, 37(5). 10.1016/j.celrep.2021.109927

Leow, L.-A., & Grahn, J. A. (2014). Neurobiology of Interval Timing. In Advances in experimental medicine and biology (Vol. 829). http://www.ncbi.nlm.nih.gov/pubmed/25358718

Levinson, S. C. (2016). Turn-taking in Human Communication - Origins and Implications for Language Processing. Trends in Cognitive Sciences, 20(1), 6–14. 10.1016/j.tics.2015.10.010

Li, J., Liao, X., Zhang, J., Wang, M., Yang, N., Zhang, J., Lv, G., Li, H., Lu, J., Ding, R., Li, X., Guang, Y., Yang, Z., Qin, H., Jin, W., Zhang, K., He, C., Jia, H., Zeng, S., … Chen, X. (2017). Primary Auditory Cortex is Required for Anticipatory Motor Response. Cerebral Cortex, 27(6), 3254–3271. 10.1093/cercor/bhx079

Li, Z., Wei, J. X., Zhang, G. W., Huang, J. J., Zingg, B., Wang, X., Tao, H. W., & Zhang, L. I. (2021). Corticostriatal control of defense behavior in mice induced by auditory looming cues. Nature Communications, 12(1), 1–13. 10.1038/s41467-021-21248-7

Matell, M. S., Meck, W. H., & Nicolelis, M. A. L. (2003). Interval timing and the encoding of signal duration by ensembles of cortical and striatal neurons. Behavioral Neuroscience, 117(4), 760–773. 10.1037/0735-7044.117.4.760

Mazzucato, L. (2022). Neural mechanisms underlying the temporal organization of naturalistic animal behavior. ELife, 11. 10.7554/elife.76577

Meck, W. H. (2006). Neuroanatomical localization of an internal clock: A functional link between mesolimbic, nigrostriatal, and mesocortical dopaminergic systems. Brain Research, 1109(1), 93–107. 10.1016/j.brainres.2006.06.031

Mita, A., Mushiake, H., Shima, K., Matsuzaka, Y., & Tanji, J. (2009). Interval time coding by neurons in the presupplementary and supplementary motor areas. Nature Neuroscience, 12(4), 502–507. 10.1038/nn.2272

Monk, K. J., Allard, S., & Hussain Shuler, M. G. (2020). Reward Timing and Its Expression by Inhibitory Interneurons in the Mouse Primary Visual Cortex. Cerebral Cortex, 30(8), 4662–4676. 10.1093/cercor/bhaa068

Namboodiri, V. M. K., Huertas, M. A., Monk, K. J., Shouval, H. Z., & Shuler, M. G. H. (2015). Visually cued action timing in the primary visual cortex. Neuron, 86(1), 319–330. 10.1016/j.neuron.2015.02.043

Narayanan, N. S., & Laubach, M. (2009). Delay activity in rodent frontal cortex during a simple reaction time task. Journal of Neurophysiology, 101(6), 2859–2871. 10.1152/jn.90615.2008

Nelken, I., Bizley, J., Shamma, S. A., Shamma, S. A., Wang, X. Q., & Wang, X. Q. (2014). Auditory cortical processing in real-world listening: The auditory system going real. Journal of Neuroscience, 34(46), 15135–15138. 10.1523/JNEUROSCI.2989-14.2014

Ning, W., Bladon, J. H., & Hasselmo, M. E. (2022). Complementary representations of time in the prefrontal cortex and hippocampus. Hippocampus, 32(8), 577–596. 10.1002/hipo.23451

Nixon, J. S., & Tomaschek, F. (2021). Prediction and error in early infant speech learning: A speech acquisition model. Cognition, 212, 104697. 10.1016/j.cognition.2021.104697

Okada, K., Matchin, W., & Hickok, G. (2018). Neural evidence for predictive coding in auditory cortex during speech production. Psychonomic Bulletin and Review, 25(1), 423–430. 10.3758/s13423-017-1284-x

Ozmeral, E. J., Eddins, A. C., Frisina, D. R., & Eddins, D. A. (2016). Large cross-sectional study of presbycusis reveals rapid progressive decline in auditory temporal acuity. Neurobiology of Aging, 43, 72–78. 10.1016/j.neurobiolaging.2015.12.024

Pendyam, S., Bravo-Rivera, C., Burgos-Robles, A., Sotres-Bayon, F., Quirk, G. J., & Nair, S. S. (2013). Fear signaling in the prelimbic-amygdala circuit: A computational modeling and recording study. Journal of Neurophysiology, 110(4), 844–861. 10.1152/jn.00961.2012

Read, H. L., Winer, J. A., & Schreiner, C. E. (2002). Functional architecture of auditory cortex. Current Opinion in Neurobiology, 12(Vi), 433–440.

Roth, B. L. (2016). DREADDs for Neuroscientists. Neuron, 89(4), 683–694. 10.1016/j.neuron.2016.01.040

Rubin, J., Ulanovsky, N., Nelken, I., & Tishby, N. (2016). The Representation of Prediction Error in Auditory Cortex. PLoS Computational Biology, 12(8), 1–28. 10.1371/journal.pcbi.1005058

Schneider, D. M., Nelson, A., & Mooney, R. (2014). A synaptic and circuit basis for corollary discharge in the auditory cortex. Nature, 513(7517), 189–194. 10.1038/nature13724

Shamash, P., Carandini, M., Harris, K., & Steinmetz, N. (2018). A tool for analyzing electrode tracks from slice histology. BioRxiv, 447995. https://www.biorxiv.org/content/10.1101/447995v1%0Ahttps://www.biorxiv.org/content/10.1101/447995v1.abstract

Smeal, R. M., Keefe, K. A., & Wilcox, K. S. (2008). Differences in excitatory transmission between thalamic and cortical afferents to single spiny efferent neurons of rat dorsal striatum. European Journal of Neuroscience, 28(10), 2041– 2052. 10.1111/j.1460-9568.2008.06505.x

Stivers, T., Enfield, N. J., Brown, P., Englert, C., Hayashi, M., Heinemann, T., Hoymann, G., Rossano, F., De Ruiter, J. P., Yoon, K. E., & Levinson, S. C. (2009). Universals and cultural variation in turn-taking in conversation. Proceedings of the National Academy of Sciences of the United States of America, 106(26), 10587–10592. 10.1073/pnas.0903616106

Sun, W., Tang, P., Liang, Y., Li, J., Feng, J., Zhang, N., Lu, D., He, J., & Chen, X. (2022). The anterior cingulate cortex directly enhances auditory cortical responses in air-puffing-facilitated flight behavior. Cell Reports, 38(10), 110506. 10.1016/j.celrep.2022.110506

Suri, H., & Rothschild, G. (2022). Enhanced Stability of Complex Sound Representations Relative to Simple Sounds in the Auditory Cortex. ENeuro, 9(4). 10.1523/ENEURO.0031-22.2022

Surlykke, A., & Moss, C. F. (2000). Echolocation behavior of big brown bats, Eptesicus fuscus, in the field and the laboratory. The Journal of the Acoustical Society of America, 108(5), 2419–2429. 10.1121/1.1315295

Taaseh, N., Yaron, A., & Nelken, I. (2011). Stimulus-specific adaptation and deviance detection in the rat auditory cortex. PLoS ONE, 6(8). 10.1371/journal.pone.0023369

Tallal, P., Miller, S., & Fitch, R. H. (1995). Neurobiological Basis of Speech: A Case for the Preeminence of Temporal Processing. The Irish Journal of Psychology, 16(3), 194–219. 10.1080/03033910.1995.10558057

Tallot, L., & Doyère, V. (2020). Neural encoding of time in the animal brain. Neuroscience and Biobehavioral Reviews, 115(June 2019), 146–163. 10.1016/j.neubiorev.2019.12.033

Town, S. M., Wood, K. C., & Bizley, J. K. (2018). Sound identity is represented robustly in auditory cortex during perceptual constancy. Nature Communications, 9(1). 10.1038/s41467-018-07237-3

Treisman, M. (1963). Temporal discrimination and the indifference interval: Implications for a model of the “internal clock”. Psychological Monographs: General and Applied, 77(13), 1–31. 10.1037/h0093864

Tremblay, P., Baroni, M., & Hasson, U. (2013). Processing of speech and non-speech sounds in the supratemporal plane: Auditory input preference does not predict sensitivity to statistical structure. NeuroImage, 66, 318–332. 10.1016/j.neuroimage.2012.10.055

Tunes, G. C., de Oliveira, E. F., Vieira, E. U. P., Caetano, M. S., Cravo, A. M., & Reyes, M. B. (2022). Time encoding migrates from prefrontal cortex to dorsal striatum during learning of a self-timed response duration task. ELife, 11, 1–19. 10.7554/eLife.65495

Ulanovsky, N., Las, L., Farkas, D., & Nelken, I. (2004). Multiple time scales of adaptation in auditory cortex neurons. Journal of Neuroscience, 24(46), 10440– 10453. 10.1523/JNEUROSCI.1905-04.2004

Ulanovsky, N., Las, L., & Nelken, I. (2003). Processing of low-probability sounds by cortical neurons. Nature Neuroscience, 6(4), 391–398. 10.1038/nn1032

Vivaldo, C. A., Lee, J., Shorkey, M. C., Keerthy, A., & Rothschild, G. (2023). Auditory cortex ensembles jointly encode sound and locomotion speed to support sound perception during movement. PLoS Biology, 21(8), e3002277. 10.1371/journal.pbio.3002277

Wang, Q., Ding, S. L., Li, Y., Royall, J., Feng, D., Lesnar, P., Graddis, N., Naeemi, M., Facer, B., Ho, A., Dolbeare, T., Blanchard, B., Dee, N., Wakeman, W., Hirokawa, K. E., Szafer, A., Sunkin, S. M., Oh, S. W., Bernard, A., … Ng, L. (2020). The Allen Mouse Brain Common Coordinate Framework: A 3D Reference Atlas. Cell, 181(4), 936–953.e20. 10.1016/j.cell.2020.04.007

Wearden, J. (2005). Origins and development of internal clock theories of psychological time. Psychologie Francaise, 50(1), 7–25.

Whitton, J. P., Hancock, K. E., & Polley, D. B. (2014). Immersive audiomotor game play enhances neural and perceptual salience of weak signals in noise. Proceedings of the National Academy of Sciences of the United States of America, 111(25). 10.1073/pnas.1322184111

Wiener, M., Lohoff, F. W., & Branch Coslett, H. (2011). Double dissociation of dopamine genes and timing in humans. Journal of Cognitive Neuroscience, 23(10), 2811– 2821. 10.1162/jocn.2011.21626

Wiener, M., Matell, M. S., & Coslett, H. B. (2011). Multiple mechanisms for temporal processing. Frontiers in Integrative Neuroscience, 5(July), 1–3. 10.3389/fnint.2011.00031

Xiong, Q., Znamenskiy, P., & Zador, A. M. (2015). Selective corticostriatal plasticity during acquisition of an auditory discrimination task. Nature, 521(7552), 348–351. 10.1038/nature14225

Yaron, A., Hershenhoren, I., & Nelken, I. (2012). Sensitivity to Complex Statistical Regularities in Rat Auditory Cortex. Neuron, 76(3), 603–615. 10.1016/j.neuron.2012.08.025

Yumoto, N., Lu, X., Henry, T. R., Miyachi, S., Nambu, A., Fukai, T., & Takada, M. (2011). A neural correlate of the processing of multi-second time intervals in primate prefrontal cortex. PLoS ONE, 6(4), 3–9. 10.1371/journal.pone.0019168

Zatorre, R. J., Belin, P., & Penhune, V. B. (2002). Structure and function of auditory cortex: Music and speech. Trends in Cognitive Sciences, 6(1), 37–46. 10.1016/S1364-6613(00)01816-7

Znamenskiy, P., & Zador, A. M. (2013). Corticostriatal neurons in auditory cortex drive decisions during auditory discrimination. Nature, 497(7450), 482–485. 10.1038/nature12077

